# Adolescent cannabinoid vapour exposure sex-dependently alters the relationship between vulnerability traits and ethanol self-administration and modifies naltrexone actions on ethanol intake in rats

**DOI:** 10.1101/2025.05.07.652663

**Authors:** Jairo S. Acosta-Vargas, Natalia de las Heras-Martínez, Alberto Marcos, Leonor Nozal, Antonio L. Crego, Marcos Ucha, Alejandro Higuera-Matas

## Abstract

**Background:** Cannabis use during adolescence is common and may predispose individuals to substance use disorders. Animal studies have explored the gateway hypothesis, but data on ethanol consumption are limited. This study aimed to investigate the potential link between adolescent cannabis exposure and ethanol self-administration, as well as the relationship between predisposing behavioural traits and ethanol consumption.

**Methods:** Adolescent rats were exposed to vapourised Δ^9^-tetrahydrocannabinol (THC) alone or with cannabidiol (CBD) at different ratios, or to a vehicle, from postnatal day (PND) 28 to 44, every other day. Behavioural assessments, including novelty and saccharin preference, goal-tracking, elevated plus maze, and ethanol self-administration (fixed and progressive ratio, punished seeking), were conducted from PND 70. Naltrexone was administered to assess its effects on ethanol intake.

**Results:** Cannabinoid exposure did not significantly affect behavioural traits or ethanol self-administration. However, sex differences emerged, with females showing a more vulnerable pattern of ethanol consumption and seeking. In THC-exposed males, a negative correlation was observed between sucrose preference and compulsive ethanol seeking, which was absent in vehicle-exposed males. In females, THC exposure disrupted the correlation between novelty preference and ethanol intake and was associated with a negative correlation between goal-tracking and compulsive seeking. Naltrexone was most effective in reducing ethanol intake in THC-exposed rats.

**Conclusions:** Adolescent cannabinoid exposure has limited effects on overall alcohol risk but may alter the psychological framework of alcohol-related behaviours and increase naltrexone potency. The observed sex differences highlight the need for personalised interventions.

## INTRODUCTION

Adolescence is a critical period of brain development during which exposure to psychoactive substances such as Δ^9^-tetrahydrocannabinol (THC) can induce long-lasting alterations in neural circuits involved in decision-making, impulse control, and reward processing (Higuera-Matas et al., 2015; Stringfield and Torregrossa, 2021). The rising prevalence of cannabis use during adolescence is particularly concerning, as it may disrupt normal brain maturation and increase vulnerability to substance use disorders, potentially acting as a gateway drug (Badiani et al., 2015; Bara et al., 2021; Fergusson et al., 2006; Kandel, 2003; Kandel et al., 2006; Lynskey and Agrawal, 2018; Secades-Villa et al., 2015). Specifically, cannabinoid exposure during this developmental window has been linked to an elevated risk of later-life substance use in rodent models (Biscaia et al., 2008; Ellgren et al., 2007; Farrelly and Vlachou, 2021; Fergusson et al., 2006; Friedman et al., 2019; Frontera et al., 2018; Higuera-Matas et al., 2008; Lecca et al., 2020; Nguyen et al., 2020; Orihuel et al., 2021; Secades-Villa et al., 2015)

In this context, our previous studies have shown that adolescent exposure to the synthetic cannabinoid CP55,940 alters neural responses to cocaine (Higuera-Matas et al., 2011) and increases cocaine self-administration in adult female, but not male, rats (Higuera-Matas et al., 2008). Additionally, adolescent THC exposure alters both the brain’s response to cocaine (Orihuel et al., 2023) and patterns of drug consumption, with adult male rats exhibiting increased cocaine intake under a progressive ratio reinforcement schedule (Orihuel et al., 2021). Other researchers have also reported a gateway effect of adolescent THC exposure on subsequent cocaine use (Friedman et al., 2019).

Few studies have examined the effects of adolescent cannabinoid exposure on alcohol consumption, and existing findings remain inconclusive. For instance, Frontera et al. (2018) found that adolescent exposure to the synthetic cannabinoid WIN 55,212-2 increased alcohol consumption in a two-bottle choice test, although this effect was measured immediately after cannabinoid exposure, without assessing long-term consequences. In contrast, (Hamidullah et al., 2021) reported that adolescent exposure to vaporized THC did not lead to increased alcohol intake in adulthood using the same paradigm. However, this study did not evaluate more pathological aspects of alcohol use, such as motivation or compulsivity (American Psychiatric Association, 2013; Deroche-Gamonet et al., 2004).

The rationale for investigating whether adolescent cannabinoid exposure may potentiate alcohol consumption and compulsive intake is grounded in evidence that such exposure alters neurotransmitter systems involved in alcohol’s reinforcing effects. Notably, it affects dopaminergic transmission (Bortolato et al., 2014; Gomes et al., 2014; Higuera-Matas et al., 2010; Pistis et al., 2004), which plays a pivotal role in the initial rewarding effects of alcohol (Vena et al., 2020), as well as glutamatergic and GABAergic signalling (Higuera-Matas et al., 2015; Orihuel et al., 2021).

Another important factor in understanding addiction vulnerability is individual variability. Certain behavioural endophenotypes, such as impulsivity, have been identified as potential risk factors (Dalley and Ersche, 2019; Toivainen et al., 2024). Our group has shown that adolescent THC exposure induces delay intolerance, particularly in females (Orihuel et al., 2021). Additional behavioural traits—including anxiety, novelty seeking, sweet preference, sign-vs. goal-tracking, and locomotor reactivity—have also been recognized as predictors of addiction susceptibility. These traits reflect distinct dimensions of behavioural and cognitive regulation, and their interplay may help explain individual differences in the risk of developing substance use disorders (Fouyssac et al., 2021). Given the predictive power of these traits and the limited research on how adolescent cannabinoid exposure might affect them concurrently, the present study aims to investigate their potential alteration.

Moreover, the route of cannabinoid administration is crucial for improving the translational relevance of animal studies. Most preclinical research has used intraperitoneal injections, which differ markedly from human routes of cannabis consumption. Recent studies have begun to use vaporization methods that more accurately mimic human use, thereby increasing ecological validity (Gutierrez et al., 2021; Hamidullah et al., 2021; Manwell et al., 2014; Moore et al., 2022, 2021; Ruiz et al., 2021; Taffe et al., 2020; Weimar et al., 2020). Similarly, incorporating cannabidiol (CBD)—a non-psychotropic cannabinoid—into experimental paradigms may better reflect real-world cannabis use. Evidence suggests that emotional and behavioural responses differ depending on whether THC is administered alone or in combination with CBD, especially when CBD concentrations are high, as in “light cannabis” products (Gabaglio et al., 2021; Pérez-Valenzuela et al., 2024). In the present study, we employ cannabis vaporization with varying THC and CBD concentrations to enhance translational relevance.

Finally, a key objective of this research is to determine whether adolescent cannabinoid exposure via vaporization not only increases vulnerability to problematic alcohol use in adulthood but also alters the efficacy of pharmacological treatments for alcohol use disorder. One such treatment is naltrexone, an opioid receptor antagonist widely used to reduce alcohol intake and prevent relapse (Anton, 2008; Bienkowski et al., 1999). Considering that THC and CBD may modulate the endogenous opioid system (Biscaia et al., 2008; Ellgren et al., 2007), it is plausible that adolescent exposure to these compounds during neurodevelopment could influence naltrexone’s therapeutic effectiveness. Exploring this interaction may yield critical insights for optimizing clinical interventions for alcohol use disorder.

## METHODS

### Animals and THC/CBD Treatment

Subjects (n=80) were the offspring of Sprague-Dawley rats (10 males and 10 females) from Charles River S.A. (Saint-Germain-sur-l’Arbresle, France), which were mated in our laboratory approximately 2 weeks after their arrival. Vaginal smears were obtained, and when the female was found to be in oestrus, she was housed with a male rat for mating. Vaginal smears were subsequently examined daily, and upon observation of spermatozoa under the microscope, the male was removed from the cage. On the day of birth (postnatal day 0), litters were sex-balanced and culled to 10±2 pups per dam so that the difference between males and females did not exceed 2 pups. The animals were weaned at 21 days of age. Different sets of animals, belonging to different litters, were randomly assigned to each experiment, minimizing litter effects. All animals were maintained at a constant temperature of 22°C ± 0.5°C with a relative humidity of 55% ± 7%, under a 12/12 h light/dark cycle (lights on at 20:00 h). Animals had ad libitum access to food (Altromin LASQdiet ROD14; Sodispan Biotech, Madrid, Spain) and tap water unless stated otherwise.

Chronic Δ^9^-tetrahydrocannabinol and cannabidiol (THCPharm, Frankfurt, Germany) treatment took place every other day from postnatal day (PND) 28 to PND 44. All solutions were prepared in glass vials under a nitrogen-saturated atmosphere and protected from light. THC (100 mg/500 µL Propylene Glycol), THC (96.97 mg) + CBD (3.03 mg) 33:1 (THChighCBDlow), THC (3.03 mg) + CBD (96.97 mg) 1:33 (THClowCBDhigh), or vehicle (500 µL Propylene Glycol) were administered on alternate days using an in-house designed vapor chamber (90 x 68 x 41 cm). Vaporization sessions lasted 30 minutes and were conducted with a Volcano system (Storz & Bickel GmbH & Co.; Tuttlingen, Germany), operating at a temperature of 230°C. Homogeneous steam distribution was facilitated by two internal small fans.

Animals were introduced into the vaporization chamber in groups of 10 males and 10 females per treatment group, in separate compartments for each sex. The vaporization sessions were conducted in the following order: first, the vehicle group (500 µL of propylene glycol); second, the pure THC group (100 mg); third, the group exposed to THC and CBD in a 33:1 ratio; and finally, the group exposed to THC and CBD in a 1:33 ratio.

Every attempt was made to minimize the pain and discomfort of the experimental animals. All procedures involving laboratory rats were conducted in accordance with the European Union legislation on the protection of animals used for scientific purposes (2010/63/EU Directive) and approved by the Ethics Board of the National University of Distance Learning and Community of Madrid (PROEX 025.1/22).

### Measurement of Rectal Temperature

Rectal temperature was measured before and after each vaporization session to evaluate the effects of cannabinoid exposure on body temperature. Baseline temperature was established by taking three consecutive measurements at 30 min, 15 min, and 0 min prior to vaporization, and the average value was used as the pre-vaporization baseline. After a 30-minute vaporization session, rectal temperature was measured again immediately. Temperature measurements were performed with a probe sensor coupled to a BIO-TK8851 digital thermometer (Bioseb, France), offering a precision of ±0.7°C. The probe was lubricated with vaseline to minimize discomfort and prevent injury to the animals. After the last vaporization session, the rats were left undisturbed until they reached adulthood when behavioural testing began.

### Determination of Cannabinoids in Plasma by LC-MS/MS

In a pilot experiment using a separate batch of rats (n=5 per group), we determined the plasma levels of THC and CBD reached with our experimental protocol. Rats were vaporized with propylene glycol, THC, or CBD for 30 minutes as described above. Immediately after this 30-minute period, the animals were anesthetized with isoflurane and decapitated. Trunk blood was collected in tubes with EDTA K3. Plasma was obtained by centrifugation at 2000 g at 4°C for 10 minutes. It was then analysed by liquid chromatography coupled to tandem mass spectrometry (LC-MS/MS) (see Supplementary Information for a detailed account of the analytic procedure). The mean amount of THC and CBD was as follows: THC [Males: 21.7 ng/mL ± 5.9 ng/mL (95% CI), Females: 22.5 ng/mL ± 3.4 ng/mL (95% CI)]. CBD [Males: 4.0 ng/mL ± 0.9 ng/mL (95% CI), Females: 2.9 ng/mL ± 1.1 ng/mL (95% CI)].

### Behavioural tests

Figure 2A depicts a timeline with all the behavioural tests performed.

### Saccharin Preference Test (Anhedonia)

At approximately PND 81, we conducted a saccharin preference test using a protocol based on previous literature (He et al., 2020). The rats were deprived of water overnight prior to the start of the test. The test began at approximately 10:00 AM and lasted for 1 hour. During the habituation phase, rats were placed in individual cages for 11 consecutive days with access to two small bottles filled with tap water (one per side) for 1 hour daily, until a stable baseline of water consumption was established. Following habituation, the saccharin preference test began, during which one of the water bottles was replaced with a freshly prepared 0.2% saccharin solution (Sigma) in tap water. To avoid positional bias, the location of the bottles was counterbalanced across sessions and animals. Saccharin preference was measured using the average consumption over the last 3 days to calculate a percentage = [Vsaccharine / (Vsaccharine + Vwater)] x 100. (V = millilitres intake)

### Elevated Plus Maze

At approximately PND 88, elevated plus maze performance was assessed (Pellow et al., 1985). The apparatus consisted of an elevated, plus-shaped maze made of black methacrylate (walls) and expanded PVC (Forex®) (floor), elevated 50 cm from the floor. The maze included two opposing closed arms (50 x 10 x 40 cm) and two open arms (50 x 10 cm). The test was conducted for a total of five minutes under 30 lux lighting conditions, following the protocol described by Higuera-Matas et al. (2009). Data were analysed as a percentage of preference using the following formula: (Time in closed arms) / (Time in open arms + Time in closed arms) *100.

### Locomotor Activity

Four days after the elevated plus maze, locomotor activity in a novel environment was also assessed using an open field (50 x 50 x 50 cm) made of Polyvinyl chloride (PVC). Rats were placed at the centre of the arena and allowed to freely explore for 60 minutes under 30 lux lighting conditions. The total distance travelled was used as a measure of locomotor activity.

### Novelty Preference Test

At approximately PND 111, preference for a novel environment versus a familiar one was assessed using a conditioned place preference (CPP) apparatus. The apparatus consisted of two compartments (30 x 50 x 50 cm) separated by a central compartment (10 x 50 x 50 cm). The compartments were divided by guillotine doors, and the walls and floor of each compartment were designed to create two distinct environments. One compartment had white walls with spherical patterns, and its floor consisted of a grid. The other compartment featured completely black walls and a floor with spherical patterns. The test began by placing the rat in the central compartment with both doors closed for 5 minutes to allow the animal to acclimate to the environment. Next, the rat was placed in one of the compartments, randomly assigned, with the door closed for 25 minutes. Finally, the rat was returned to the central compartment with the doors open, and its preference for each compartment was recorded for 15 minutes. Data were analysed using the following formula: (novel compartment time) / (novel compartment time + familiar compartment time) x100.

### Pavlovian Conditioned Approach

At approximately PND 124, the Pavlovian Conditioned Approach (PCA) procedure was initiated. The day before the test, rats were food-restricted for 12 hours to ensure motivation during the task and continued to be so until the end of this procedure. The first session consisted of a 60-minute habituation period in the Skinner boxes, during which the levers remained retracted, and the animals were exposed to the chamber environment without receiving pellets, following the protocol described in Orihuel et al. (2021a). The subsequent 9 daily training sessions consisted of 25 trials per session (30 minutes), during which a pellet was dispensed into the magazine according to a variable interval 60-second (VI-60) schedule of reinforcement. A lever (positioned on the right or left, counterbalanced between animals) was extended for 8 seconds (8 s) immediately before pellet delivery and retracted right after. The other lever remained present throughout the entire session and served as a control to measure general locomotor activity, as neither lever had programmed contingencies. During these 9 sessions, the rats were subjected to a food-restriction protocol to maintain their body weight between 90% and 95% of their theoretical free-feeding weight.

During PCA sessions, the interaction with the conditioned stimulus (CS)+ lever (CS+ LPs), presses on the inactive lever in between each CS+ (ILPs) or during the CS+ (CS+ ILPs), and Head Entries (HE) in between each CS+ presentation and during the CS+ (CS+ HE) were recorded. The main PCA index was calculated each day as the mean of three other indexes:

1. Response bias, i.e., the ratio of the total number of LP and HE for a session during the CS+ presentations: (CS+LPs - CS+HEs) / (CS+LPs + CS+HEs)
2. Latency score, i.e., average latency to perform a LP or HE during the 8 s of CS+ presentation: (Mean HEs latency - Mean LP latency)/8
3. Probability difference, i.e., the probability of performing a HE during the CS+ presentations minus the probability of performing a LP during the CS+ presentation: P(LP) - P(HE)

Every index score ranged from −1 (absolute sign-tracking) to 1 (absolute goal-tracking). Animals with a PCA score higher than 0.5 were classified as goal-trackers, whereas animals with a PCA score lower than −0.5 were categorized as sign-trackers. Scores from 0.5 to −0.5 are considered an intermediate phenotype with no clear preference bias.

### Video Tracking

Animal behaviour was recorded by an overhead video camera and analysed using the software ANY-maze Tracking System v.6.32 (Stoelting Co.).

### Acquisition of Ethanol self-administration, progressive ratio performance and study of punished seeking behaviour

On PND 138, we initiated the ethanol self-administration protocol as described (see Supplementary Information for a more detailed description of all the phases). The operant ethanol self-administration protocol was designed to assess the development of ethanol-seeking and -taking behaviours, the motivation to obtain ethanol, and the specificity of the response to the active lever and the compulsivity in ethanol seeking, and it was structured into the following seven distinct phases:

#### Acquisition

Acquisition was conducted without prior saccharin masking, following the protocol described by the Heillig lab (Augier et al., 2018; Domi et al., 2021). Sessions lasted 30 minutes and began with the extension of both the active and inactive levers to signal the availability of the drug. Pressing the active lever triggered the pump, which dispensed an ethanol dose (100 μL at 15% v/v) into the drinking receptacle, allowing the rat to voluntarily consume it. The inactive lever had no programmed contingencies and served as a control for non-specific motor activity.

A total of 24 sessions (one per day) were conducted under a fixed ratio (FR1) schedule, in which the rat had to press the active lever once to receive a single ethanol delivery (100 μL at 15% v/v). After these 24 sessions, the schedule was shifted to FR2 for an additional 14 sessions, where the rat had to press the active lever twice to obtain a single delivery of ethanol. The basic characteristics of these sessions under the FR2 schedule remained the same for subsequent phases of the operant protocol.

#### Progressive ratio

To assess motivation for ethanol intake, a progressive ratio (PR) schedule was conducted. PR conditions were identical to acquisition phase but with an increased response requirement per ethanol reinforcer (increasing behavioural cost) according to the following scale: 1, 2, 4, 6, 9, 12, 15, 20, 25, 32, 40, 50, 62, 77, 95, 118, 145, 178, 219, 268. The self-administration session ended after 30 min. This PR series was derived from the following equation: = [5e ^(injection^ ^number^ ^X^ ^0.2)^ - 5]. Following the protocol described by (Richardson and Roberts, 1996). The breaking point was defined as the last lever press requirement reached that resulted in ethanol administration

#### Stabilization

After progressive ratio (PR) schedule, animals were allowed to recover lever pressing for ethanol self-administration behaviour over seven sessions in absence of PR under a fixed ratio 2 (FR2) schedule.

#### Compulsive Consumption

After stabilization sessions, compulsivity was operationalized as responding for ethanol when its delivery was associated with a foot-shock punishment. Briefly, animals underwent twenty-two sessions in which every two lever presses (FR2) were paired with a 0.5-s footshock, contingent on the subsequent delivery (3 seconds later) of 100 μL of 15% ethanol in water, which was deposited by activation of the pump for 4.5 s into the drinking receptacle. The stimulus light above the active lever was illuminated concurrently, both lasting for the same duration. The intensity of the electric shock used as punishment was 0.2 mA, consistent with previous findings showing that thin intensity reveals the compulsive phenotype (Domi et al., 2021).

#### Stabilization

After footshock punishment, male and female rats were allowed to recover lever pressing for ethanol in the absence of punishment. Five sessions under the FR2 schedule were conducted to return to baseline levels of ethanol self-administration. To habituate the animals to the injection procedure and reduce stress before the subsequent naltrexone administration phase, they received a single daily intraperitoneal (i.p.) injection of sterile physiological saline (1 mL/kg) 30 minutes before sessions 3, 4, and 5.

#### Naltrexone Administration

Male and female rats received a single i.p. administration of 0.3 mg/kg naltrexone, 30 minutes before the start of the self-administration ethanol session under the FR2 schedule (i.e., 2 responses), as described by (Augier et al., 2017). Naltrexone was dissolved in sterile physiological saline and administered in a volume of 1 mL/kg.

#### Statistical Analysis

IBM SPSS Statistics v.27.0 was used to analyse the data. All results are expressed as the mean ± SEM (Standard Error of the Mean). Mixed ANOVAs were conducted to evaluate the effects of within-subject factors (sessions and lever type: active vs. inactive) and between-subject factors (sex and treatment), as well as their interactions. Univariate two-way ANOVAs were used to analyse data with one dependent variable and between-subject factors (sex and treatment) in cases where no repeated measures were involved. Greenhouse-Geisser corrections were applied to F-values when the assumption of sphericity was violated. Significant interactions and main effects were followed up with simple effects analysis or pairwise comparisons, with Bonferroni corrections applied when necessary. F-values, effect sizes (partial eta squared, ηp^2^), and degrees of freedom were reported wherever appropriate, with statistical significance set at p < 0.05. Pearson bivariate correlations were computed between behavioural predictors and different indices of ethanol self-administration.

## Results

### Hypothermia Induced by Adolescent Cannabinoid Vapor Exposure

Exposure to vaporized cannabinoids decreased rectal temperature, an effect that was partially mitigated by the presence of CBD in the mixture. Analysis of post-vaporization group differences indicated that the strongest hypothermic effect was observed in the THC group, followed by the THChighCBDlow group, and then the THClowCBDhigh group (Figure 1A). In the last two vaporization sessions, a similar pattern of results was obtained, except for the stronger hypothermic effect observed in the THChighCBDlow group, which differed from the THClowCBDhigh group after vaporization (Figure 1B). Tables S1-S6 provide all the statistical data of the main effects and pairwise comparisons performed. Figures S1 shows the data for each sex. For a more detailed account of the rectal temperature results, see the corresponding section in the supplementary results.

**Figure 1.**
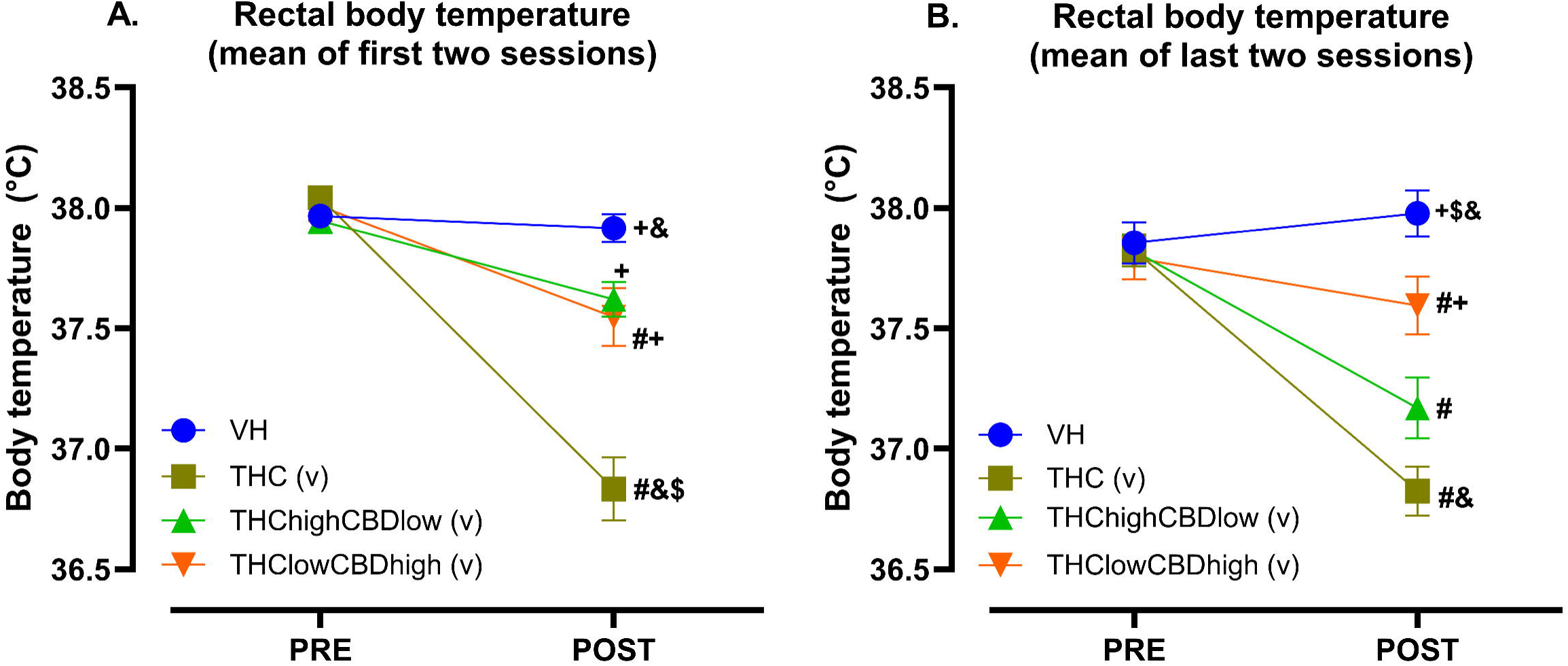
Effects of cannabinoid vaporisation on rectal temperature. (A) The mean rectal temperature from the first two sessions (pre- and post-vaporisation). Within-subject effects were not observed only in the vehicle (VH) group indicating stable rectal temperature in this group. However, all cannabinoid-treated groups exhibited a significant reduction in rectal temperature following vaporisation, THC group (p < 0.001), followed by THChighCBDlow (p < 0.001) and THClowCBDhigh (p = 0.001), as indicated in the pairwise comparisons. Between-subject effects post-vaporisation, THC exhibited the most pronounced hypothermic effect, showing a significant reduction in rectal temperature compared to the VH, THChighCBDlow, and THClowCBDhigh groups. In contrast, the THChighCBDlow group showed a significant reduction compared to THC but no from VH and THClowCBDhigh. (B)The mean rectal temperature from the last two sessions (pre- and post-vaporisation). Within-subject effects were not observed the vehicle (VH) group, indicating stable rectal temperature across conditions. However, all cannabinoid-treated groups exhibited a significant reduction in rectal temperature following vaporisation. This hypothermic effect was strongest in the THC group (p < 0.001), followed by THChighCBDlow (p < 0.001) and THClowCBDhigh (p = 0.033), as indicated in the pairwise comparisons. Between-subject effects post-vaporisation, THC induced hypothermia remained significantly different from VH and THClowCBDhigh, but not from THChighCBDlow. The THClowCBDhigh group only exhibited significantly different from VH. Symbols denote significant differences (*p < 0.05), from the vehicle (VH) is indicated by ^#^, THC is indicated by ^+^, THChighCBDlow is indicated by ^$^ and THClowCBDhigh is indicated by ^&^. Significant effect of the within-subjects factor (pre-post vaporisation effect of rectal temperature) is denoted by ‘v’. All data are expressed as mean ± S.E.M.

### Behavioural Tests

#### Saccharine Preference Test

This test was conducted to evaluate potential diminished preference for rewards (anhedonia) induced by cannabinoid exposure during adolescence. Anhedonia was measured as the preference for a sweet solution (saccharine) over water across 10 sessions. When we analysed the mean of the last three sessions, there was a trend for an effect of treatment (F(3, 70) = 2.531, p = 0.064, ηp^2^ = 0.098) and no significant effect of sex (F(1, 70) = 0.829, p = 0.366, ηp^2^ = 0.012) or the sex x treatment interaction (F(3, 70) = 0.014, p = 0.998, ηp^2^ = 0.001) (Figure 2B).

**Figure 2.**
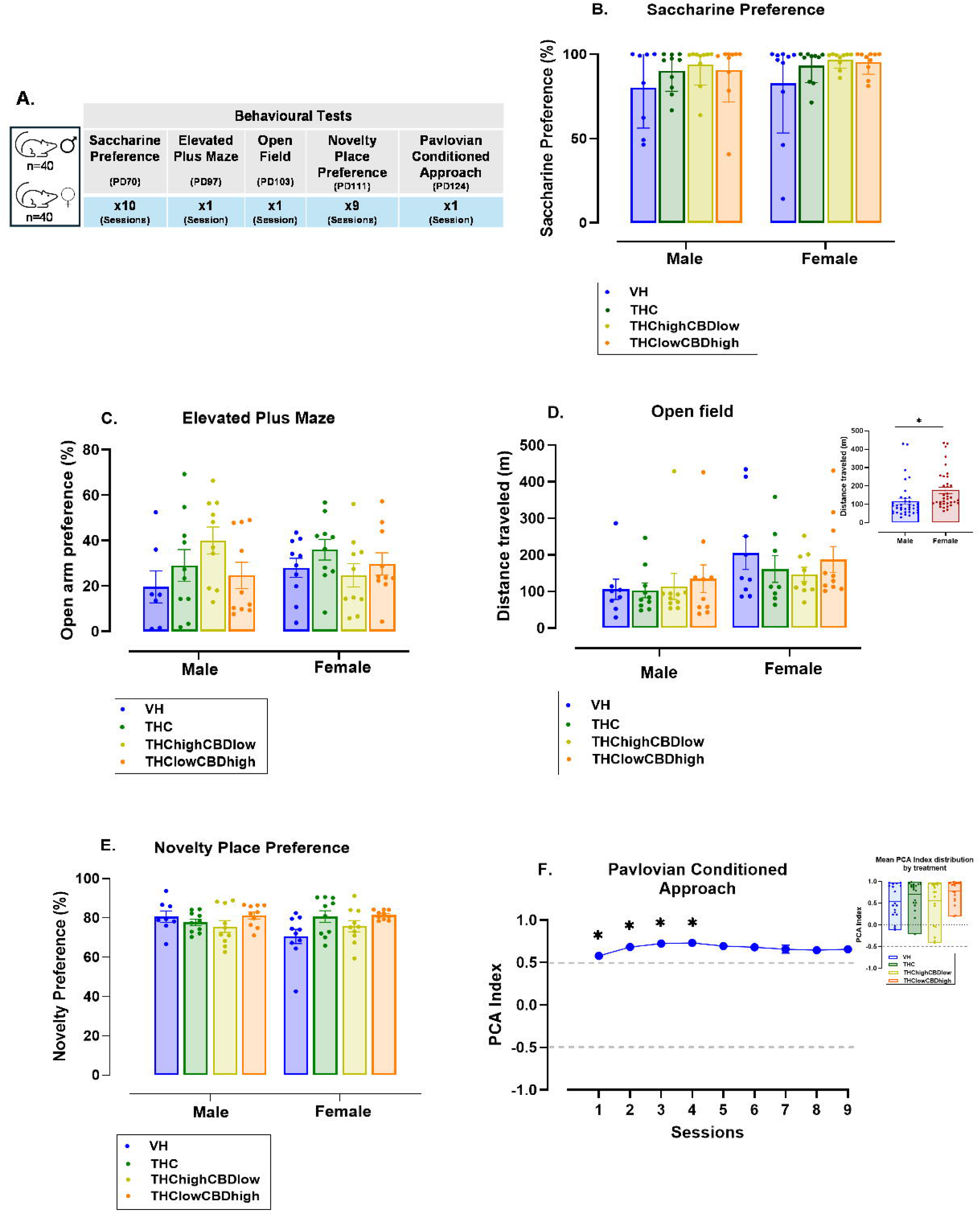
Behaviour test characterization. (A) Sequence and duration of each behavioural test. (B) Saccharine Preference Test. The percentage of saccharine preference is shown for male and female subjects across treatment groups (± S.E.M.), no significant differences were observed, indicating that adolescent cannabinoid exposure did not affect hedonic behaviour in adulthood. (C) Elevated Plus Maze Test. The mean percentage of time spent in the closed arms is presented for male and female subjects across treatment groups (± S.E.M.). The results suggest that adolescent cannabinoid exposure did not significantly influence anxiety-related behaviour in adulthood. (D) Open Field Test. The distance travelled by males and females during the open field test is displayed (± S.E.M.). Female subjects exhibited higher locomotor activity compared to males, independent of the treatment group. (E) Novelty Preference Test. The percentage of novelty preference is depicted for male and female subjects across treatment groups (± S.E.M.). No significant differences were observed, suggesting that neither sex nor adolescent cannabinoid exposure affected exploratory behaviour in adulthood. (F) Pavlovian Conditioned Approach Test. The mean PCA index across nine sessions is shown, illustrating changes in conditioned responses over time (± S.E.M.). A significant effect of sessions was observed, indicating an initial increase in approach behaviour that stabilized after session 4. However, neither sex nor treatment significantly influenced the conditioned response. (see insert). Post hoc analysis (Bonferroni correction) revealed that responses in sessions 2, 3, and 4 were significantly different from session 1 (*p < 0.05).

#### Elevated Plus Maze

No significant differences were found in the percentage of time spent in the open arms due to the effects of sex (F(1, 70) = 0.234, p = 0.630, ηp^2^ = 0.003) or treatment (F(3, 70) = 1.414, p = 0.246, ηp^2^ = 0.057). Additionally, no significant interaction effects were observed between sex and treatment (F(3, 70) = 2.232, p = 0.092, ηp^2^ = 0.087). These results suggest that adolescent cannabinoid exposure did not significantly alter anxiety-related behaviour in adulthood (Figure 2C).

#### Open Field Maze

Significant differences were found due to the effect of the sex factor (F(1, 66) = 6.579, p = 0.013, ηp^2^ = 0.091), with females exhibiting significantly higher locomotor activity compared to males (mean difference = 60.909, p = 0.013). However, no significant differences were observed due to the treatment factor (F(3, 66) = 0.461, p = 0.710, ηp^2^ = 0.021) or the sex x treatment interaction (F(3, 66) = 0.333, p = 0.801, ηp^2^ = 0.015) (Figure 2D).

#### Novelty Preference Test

This test was conducted to assess exploratory behaviour and preference for novel environments. No significant differences were found due to the effects of sex (F(1, 70) = 0.859, p = 0.357, ηp^2^ = 0.012) or treatment (F(3, 70) = 2.331, p = 0.082, ηp^2^ = 0.091). Additionally, no significant sex x treatment interaction effects were observed (F(3, 70) = 2.310, p = 0.084, ηp^2^ = 0.090) (Figure 2E).

#### Pavlovian Conditioned Approach (PCA)

This test was conducted to evaluate goal- and sign-tracking behaviour, which reflects individual differences in conditioned responses to reward-associated cues, often linked to addiction vulnerability. Significant differences were found due to the effect of the sessions factor (F(3.051, 213.54) = 3.311, p = 0.02, ηp^2^ = 0.045), indicating that responses improved or changed across sessions (Figure 2F). Post-hoc analysis with Bonferroni correction revealed that these differences were only present during the first four sessions compared to the first session (p < 0.05), after which responses stabilized. However, no significant differences were observed due to the interaction of sessions x sex (F(3.051, 213.54) = 2.158, p = 0.93, ηp^2^ = 0.013) or sessions x sex x treatment (F(9.152, 213.54) = 0.276, p = 0.981, ηp^2^ = 0.012). Additionally, the mean performance of the last three sessions was analysed to assess potential effects of sex and treatment. No significant main effects or interactions were found. Specifically, no differences were observed for sex (F(1, 70) = 1.162, p = 0.285, ηp^2^ = 0.016), treatment (F(3, 70) = 1.748, p = 0.165, ηp^2^ = 0.070), or the interaction of sex × treatment (F(3, 70) = 0.448, p = 0.719, ηp^2^ = 0.019). These findings suggest that while responses varied across sessions, neither sex nor treatment significantly influenced conditioned approach behaviour, even when focusing on the mean of the last three sessions (Figure 2F).

### Ethanol Self-Administration

#### Acquisition – Fixed Ratio 1 (FR1)

During the FR1 phase, animals progressively learned to press the active lever to obtain a dose of ethanol, as evidenced by a significant main effect of sessions (F(13.026, 898.771) = 4.837, p < 0.001, ηp^2^ = 0.066), lever (F(1, 69) = 93.425, p < 0.001, ηp^2^ = 0.575), and the interaction of sessions × lever (F(11.812, 815.001) = 16.107, p < 0.001, ηp^2^ = 0.189). This result confirms that ethanol self-administration behaviour increased over time as sessions progressed (Figure 3A). The interaction between sessions × treatment was also significant (F(39.077, 898.771) = 1.606, p = 0.012, ηp^2^ = 0.065), suggesting that acquisition rates varied between treatment groups, but this effect was limited to two sessions and was considered spurious (see Supplementary Information).

**Figure 3.**
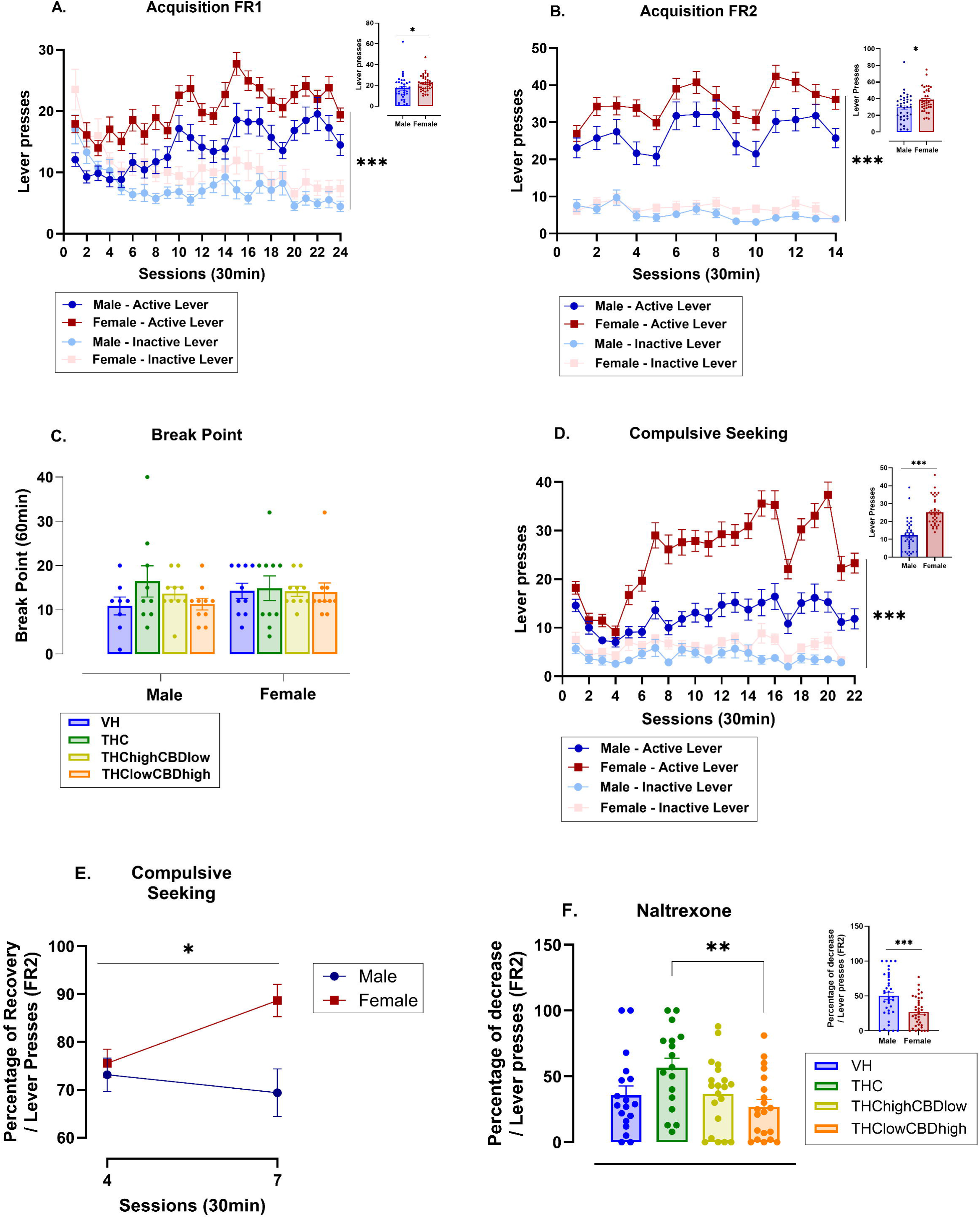
Ethanol self-administration, motivation and compulsivity behaviour, and the effectiveness of naltrexone. (A) Discrimination between active and inactive lever presses during the acquisition phase (FR1) across 24 sessions (± S.E.M.). Females consistently pressed the active lever more than males throughout all sessions. (B) Discrimination between active and inactive lever presses during the maintenance phase (FR2) across 14 sessions (± S.E.M.). Similar trends were observed, with females exhibiting a higher frequency of active lever presses compared to males. (C) Breaking Points achieved by sex between treatments (± S.E.M.). There was no significant effect of the treatment; however, a notable effect of sex was observed, as females reached a higher ratio of active lever presses in the progressive ratio compared to males. (D) Active lever presses by sex during the compulsive seeking across 22 sessions (± S.E.M.). Females displayed greater compulsive behaviour than males during the compulsivity test, even when receiving punishment for pressing the active lever in their pursuit of ethanol. (E) Mean percentage change in recovery by sex during the compulsivity test (± S.E.M.), the females recovered a greater percentage of their ethanol-seeking behaviour after receiving punishment for active lever pressing compared to males, highlighting their more compulsive tendencies. (F) Mean percentage change in the decrease of active lever presses by sex following naltrexone administration (± S.E.M.). The group treated with THC exhibited a greater percentage reduction in lever presses compared to the THClowCBDhigh group. Additionally, males generally showed a more pronounced decrease in ethanol-seeking behaviour than females after administration of naltrexone. Significance symbols are defined as follows: * indicates p < 0.05, ** indicates p < 0.01, and *** indicates p < 0.001.

We also observed a main effect of sex, with females exhibiting significantly more lever presses compared to males (F(1, 69) = 14.138, p < 0.001, ηp^2^ = 0.170). These results suggest that animals treated with cannabinoid vapor during adolescence did not show different operant behaviour for ethanol self-administration compared to the VH group, and that, as a group, females self-administered more ethanol than males.

#### Acquisition – Fixed Ratio 2 (FR2)

During the FR2 phase, a significant main effect of lever was observed (F(1, 69) = 277.028, p < 0.001, ηp^2^ = 0.801), indicating a strong preference for the active lever over the inactive lever (Figure S3B), thus demonstrating that the animals had successfully acquired the operant behaviour. Additionally, a significant interaction between lever and sex (F(1, 69) = 5.339, p = 0.024, ηp^2^ = 0.072) suggested that females exhibited slightly higher active lever-pressing activity compared to males, showing a marked sexual difference. However, there was no significant interaction between sessions and treatment (F(39, 897) = 1.244, p = 0.148, ηp^2^ = 0.051), lever × treatment (F(3, 69) = 0.718, p = 0.544, ηp^2^ = 0.030), or the three-way interaction of lever × sex × group (F(3, 69) = 1.075, p = 0.366, ηp^2^ = 0.045).

The analysis of the mean number of active lever presses during the last three sessions revealed a statistically significant between-subjects effect for the sex factor (F(1, 69) = 6.874, p = 0.011, ηp^2^ = 0.091). Post-hoc comparisons with Bonferroni corrections showed that males pressed the active lever less frequently than females (mean difference = −8.949, SE = 3.413, p = 0.011, 95% CI [−15.757, −2.140]) (Figure 2B).

These findings underscore that while the preference for the active lever over the inactive lever increased significantly across sessions, this trend was consistent across sexes, reflecting robust acquisition of the operant behaviour under the FR2 schedule. At the end of FR2 training, females showed more robust responding for ethanol than males. The absence of significant interactions involving treatment suggests that cannabinoid exposure did not significantly alter the maintenance of ethanol self-administration assessed in this phase.

#### Progressive Ratio

Motivation for ethanol consumption was assessed using a progressive ratio (PR) schedule. In this schedule, the number of active lever presses required to receive each subsequent ethanol dose increased progressively, making the task more demanding as the session advanced. The last lever press requirement that resulted in an ethanol dose administration is defined as the breakpoint. The analysis of the breakpoint achieved showed that there were no significant differences due to the effect of sex (F(1, 69) = 0.721, p = 0.399, ηp^2^ = 0.010), treatment (F(3, 69) = 0.926, p = 0.433, ηp^2^ = 0.039), or the interaction between sex and treatment (F(3, 69) = 0.555, p = 0.646, ηp^2^ = 0.024) (Figure 3C). Thus, neither sex nor treatment had any significant impact on operant behaviour related to the motivation to consume ethanol. There were no significant effects for lever presses in this phase (Figure S4). See Supplementary Information for the detailed statistical analysis.

#### Compulsive Ethanol Seeking

In the compulsive ethanol seeking phase, animals were subjected to a fixed ratio 2 (FR2) schedule, where pressing the active lever first delivered a 0.2 mA footshock to punish the seeking behaviour, followed by activation of the pump that dispensed a dose of ethanol, which the rat then consumed. Significant differences were observed for the main effect of lever (F(1, 67) = 262.079, p < 0.001, ηp^2^ = 0.796), confirming a strong preference for the active lever despite the punishment. The three-way interaction of lever × sessions × sex showed significant effects (F(11.468, 779.628) = 2.334, p = 0.008, ηp^2^ = 0.033), revealing a gradual increase in lever pressing in the absence of ethanol reward (Figure 3D). Post-hoc comparisons with Bonferroni corrections showed that males and females pressed the active lever at similar levels during sessions 1 and 2, but from session 3 onward, females pressed the active lever more than males across the remaining sessions. There was no significant effect of the lever x sessions x treatment interaction (F(34.403, 768.338) = 1.040, p = 0.407, ηp^2^ = 0.045), nor was there any effect of the lever x sessions x sex x treatment interaction (F(34.403, 768.338) = 1.017, p = 0.442, ηp^2^ = 0.044) (Figure S6C).

We also calculated the percentage decrease in the operant response (active lever presses) after the onset of electric shocks as punishment, relative to the average of the last three sessions of the stabilization phase (without electric shocks). After session 4, the rats reached their maximum decrease, and from session 5 to 7, they achieved their peak recovery. We observed significant differences due to the interaction between the factors sessions x sex (F(1, 67) = 4.764, p = 0.033, ηp^2^ = 0.066) (Figure 3E), with females exhibiting more pronounced compulsive behaviour compared to males in terms of ethanol self-administration. We did not observe significant differences between the session x treatment factor (F(3.000, 67.000) = 1.225, p = 0.307, ηp^2^ = 0.052) or the interaction between the factors sessions x sex x treatment (F(3.000, 67.000) = 0.539, p = 0.657, ηp^2^ = 0.024).

Importantly, there was a significant correlation between active lever presses (and rewards) in this phase and head entries to the ethanol-containing receptacle (Figure S6 A and B and Table S7), confirming that rats were consuming the ethanol during this phase.

#### Naltrexone Administration

After completing the ethanol self-administration phase with punishment (footshock punishment), animals underwent five additional operant self-administration sessions without punishment. This was done to extinguish the association between ethanol-seeking behaviour and the punishment previously received. Following this stabilization period, the efficacy of naltrexone in reducing ethanol-seeking behaviour was assessed. The percentage reduction in ethanol self-administration following naltrexone administration (given 30 minutes before testing) was calculated as the ratio of total active lever presses after naltrexone administration to the sum of the mean active lever presses from the last three sessions without punishment and total active lever presses after naltrexone administration, multiplied by 100 to express the reduction as a percentage.

Significant differences were observed in the percentage reduction due to the factor sex (F(1, 67) = 16.381, p < 0.001, ηp^2^ = 0.196), with females showing a smaller reduction in lever pressing compared to males (mean difference = 23.677, p < 0.001, 95% CI [12.000, 35.354]), indicating a lower effectiveness of naltrexone in females. Significant differences were also found due to the factor treatment (F(3, 67) = 4.028, p = 0.011, ηp^2^ = 0.153), and the sessions x treatment interaction (F(3, 61) = 2.987, p = 0.038, ηp^2^ = 0.128), where the THC treatment group exhibited the highest percentage reduction in lever pressing (mean difference compared to THClow_CBDhigh = 28.642, p = 0.006, 95% CI [6.003, 51.280]), highlighting a stronger naltrexone effect in reducing ethanol-seeking behaviour in this group (Figure 3F). However, no significant differences were found for the interaction of the factors sessions x sex x treatment (F(3, 61) = 0.654, p = 0.583, ηp^2^ = 0.031).

### Correlation Between Behavioural Characterization and Ethanol Self-Administration Phase

To investigate the potential of various behavioural parameters as predictors of ethanol consumption in adulthood, including compulsive consumption, Pearson correlation analysis were conducted. The results revealed significant correlations between these behavioural parameters and ethanol consumption, although only one significant correlation was observed in males, in contrast to six significant correlations found in females. Figures 4A and 4B present heatmaps of the Pearson correlation coefficients for all four treatment groups in males and females. Table S8 contains a summarises these data.

**Figure 4.**
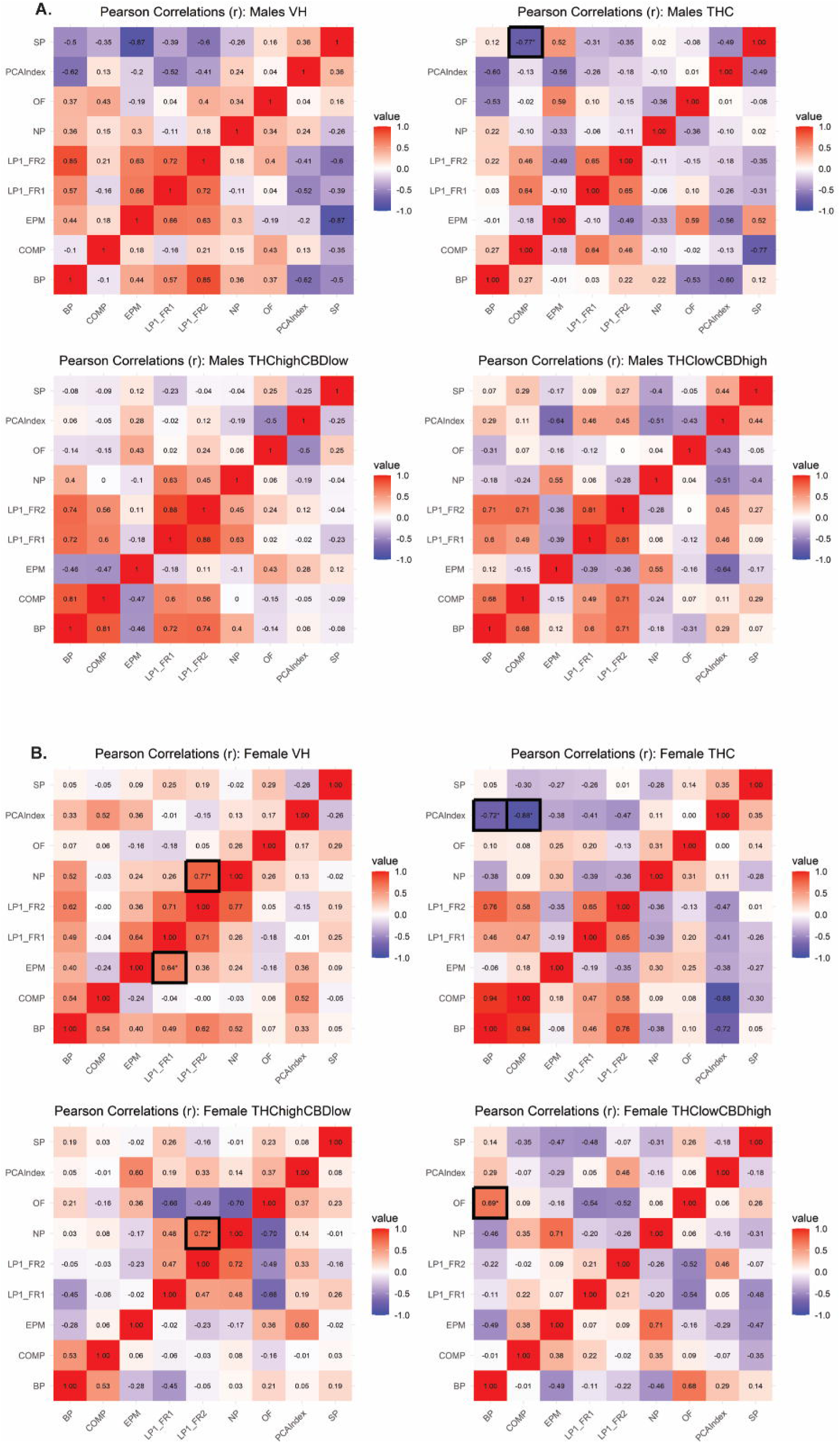
Heatmap of Pearson Correlations by treatment groups in males. (A) Pearson correlation coefficients (r) in the four treatment groups (VH, THC, THChighCBDlow, and THClowCBDhigh) for the males. (B) Pearson correlation coefficients (r) in the four treatment groups (VH, THC, THChighCBDlow, and THClowCBDhigh) for the females.

### Males

#### Saccharine Preference (SP) and Compulsivity (COMP)

A significant negative correlation was observed between saccharine preference and compulsivity in males treated with THC (r = −0.769, p = 0.015). This suggests that higher preference for saccharine during adolescence is associated with lower compulsivity in ethanol consumption during adulthood in this group.

### Females

#### Elevated Plus Maze (EPM) and Lever Presses – FR1

In females treated with vehicle (VH), a significant positive correlation was found between time spent in the open arms of the EPM and lever presses under an FR1 schedule (r = 0.639, p = 0.047).

#### Novelty Preference and Lever Presses – FR2

In females treated with vehicle (VH), a significant positive correlation was found between novelty preference and lever presses under an FR1 schedule (r = 0.766, p = 0.010).

#### PCA Index and Breaking Point

In females treated with THC, a significant negative correlation was found between PCA Index and breaking point in females treated with THC (r = −0.717, p = 0.020), indicating that higher PCA Index scores (goal-tracking behaviour) are associated with lower motivation to obtain ethanol in adulthood.

#### PCA Index and Compulsivity

A strong negative correlation was observed between PCA Index and compulsive drinking behaviour in females treated with THC (r = −0.880, p = 0.004).

#### Novelty Preference and Lever Presses – FR2

In females treated with THChighCBDlow, a significant positive correlation was observed between novelty preference and lever presses under an FR2 schedule (r = 0.724, p = 0.018).

#### Open Field Test and Breaking Point

In females treated with THClowCBDhigh, a significant positive correlation was found between activity in the open field test and breaking point (r = 0.681, p = 0.030).

## Discussion

This study demonstrates that adolescent exposure to vaporized cannabinoids did not significantly impact saccharine preference, anxiety-related behaviours, novelty preference, or Pavlovian conditioned approach, all of which have been linked to various aspects of drug self-administration or compulsive seeking (Anderson and Spear, 2011; Belin et al., 2011; Dilleen et al., 2012; Fouyssac et al., 2021; Gosnell et al., 1998). However, it did alter the relationship between some of these behavioural traits and certain aspects of ethanol self-administration. Furthermore, the presence of CBD in the cannabinoid mixture modulated some of these effects. Notably, the pattern of results suggests a heightened suppression of ethanol intake by naltrexone in THC-exposed rats.

### Sex Differences in Ethanol Self-Administration

Sex differences were prominent across nearly all ethanol self-administration measures, with females demonstrating higher ethanol consumption than males under both easy access and punished-seeking conditions. Additionally, naltrexone exhibited lower potency in females. This aligns with previous studies showing that females often exhibit increased ethanol self-administration (Flores-Bonilla and Richardson, 2020; Hilderbrand and Lasek, 2018; Maddern et al., 2024; Salazar and Centanni, 2024; Towers et al., 2022), a pattern that is also reflected in operant paradigms when adjusted for weight (Randall et al., 2017; Toivainen et al., 2024).

Moreover, hormone fluctuations during the oestrus cycle do not appear to influence ethanol intake in operant self-administration models (Bertholomey and Torregrossa, 2019; Randall et al., 2017) However, studies have shown that oestradiol replacement in ovariectomized rats increases ethanol consumption compared to vehicle-treated ovariectomized or intact females (Bertholomey and Torregrossa, 2019). These sex differences in ethanol consumption may be due to a heightened dopaminergic response to ethanol in females, as previous research has indicated greater dopamine release in response to ethanol in females compared to males (Blanchard and Glick, 1995) Additionally, differences in the glutamate, GABA, opioid, and endocannabinoid systems, which exhibit significant sex-based variability (Higuera-Matas et al., 2012; Lafta et al., 2024; Viveros et al., 2012), may also contribute to these patterns.

### Thermal Responses and Cannabinoid Interactions

Rats exposed to vaporized THC exhibited the well-documented hypothermic response to THC (Moore et al., 2021; Nguyen et al., 2020, 2016). However, combining THC with CBD attenuated this effect. Previous studies have shown similar results with THC+CBD mixtures, though the effect on hypothermia was more pronounced in females when using a 1:4 THC:CBD ratio (Javadi-Paydar et al., 2018). Our findings align with those of Todd and Arnold (2016), who reported that CBD mitigates THC-induced hypothermia at a 1:1 ratio. These effects may be mediated by CB1 receptors in hypothalamic regions, such as the medial preoptic nucleus, which are involved in thermoregulation (Todd and Arnold, 2016).

### Long-Term Effects of Adolescent Cannabinoid Exposure on Drug-Related Behaviours

Adolescent cannabinoid exposure did not significantly alter behaviours that are predictive of compulsive drug use, such as saccharine preference, anxiety-related behaviours, or Pavlovian conditioned approach. While the literature on the long-term effects of adolescent cannabinoid exposure is extensive (Farrelly and Vlachou, 2021; Higuera-Matas et al., 2015; Martín-Cuevas et al., 2023; Rubino and Parolaro, 2016; Zamberletti and Rubino, 2022) studies specifically examining the effects of vaporized cannabinoids during adolescence are more limited (Freels et al., 2024; Gutierrez et al., 2024; Nguyen et al., 2020). This study provides the first report of the long-term effects of vaporized THC and THC+CBD exposure on these behaviours in rodents. Despite achieving physiologically relevant THC plasma levels (21.7 ng/ml in males and 22.5 ng/ml in females), we found no long-term effects on the behaviours assessed. However, the doses used here were lower than those reported in other studies (Nguyen et al., 2020), suggesting that future research using higher doses may reveal dose-dependent effects.

### Ethanol Self-Administration and the Gateway Hypothesis

This study sought to assess the potential “gateway” effect of adolescent THC vapor exposure on ethanol self-administration, including behaviours indicative of alcohol use disorder, such as increased motivation and compulsive drug seeking. Previous studies focusing on the effects of cannabinoid exposure on ethanol intake during adolescence have either examined free ethanol consumption (Frontera et al., 2018) or voluntary ethanol consumption in adolescent rats without assessing motivations related to alcohol use disorder (Hamidullah et al., 2021). Contrary to expectations, exposure to cannabinoids during adolescence did not significantly impact operant ethanol self-administration, except for some isolated effects during the FR1 phase. These results align with previous reports that found no effects of adolescent THC exposure on free-access ethanol intake(Hamidullah et al., 2021). Further studies using higher doses or ascending dose schedules (e.g., Moreno-Fernández et al., 2025) are needed to explore the potential gateway effect observed with other substances, such as opiates (Maddern et al., 2024; Nguyen et al., 2020).

### Modulatory Effects of THC and CBD on Ethanol-Related Behaviours

While cannabinoid exposure did not result in significant long-term effects on ethanol self-administration, it did alter the relationship between certain behavioural traits and ethanol consumption. In male rats exposed to THC vapor, a significant negative correlation emerged between sucrose preference and compulsive ethanol seeking, a relationship that was absent in vehicle-exposed controls. This suggests that THC exposure may shift the relationship between reward-seeking and avoidance of aversive stimuli, favouring approach behaviour. In contrast, in females, THC exposure disrupted the typical correlation between novelty preference and ethanol self-administration, which was observed in vehicle-exposed controls (Toivainen et al., 2024). Notably, a high THC and low CBD mixture restored the correlation between novelty preference and ethanol intake, suggesting a modulating effect of CBD. These findings highlight how cannabinoid exposure during adolescence can sex-dependently alter the predictive value of behavioural traits associated with ethanol consumption.

## Conclusion

This study provides compelling evidence that exposure to THC vapor during adolescence enhances the potency of naltrexone in reducing ethanol intake. Additionally, cannabinoids—particularly depending on their THC:CBD ratio—disrupt the linear relationship between psychological predictors of drug use and different aspects of alcohol use disorder in a rodent self-administration model. These findings have potential clinical implications, suggesting that clinicians should consider a patient’s history of cannabis use when diagnosing and treating alcohol use disorder, as cannabinoid exposure may modify both the diagnostic process and therapeutic responses (e.g., to naltrexone). Furthermore, the observed sex differences in ethanol-related behaviours should be considered when designing personalized preventive and therapeutic interventions

### Ethics statement

The authors do not have any conflict of interest to disclose. All experimental procedures were previously approved by the Ethics Committee of UNED and the Autonomous Community of Madrid (PROEX 025.1/22). The animals were maintained and handled according to the European Union Laboratory Animal Care Rules (EU Directive 2010/63/EU) and the “Principles of Laboratory Animal Care” were strictly followed. This manuscript has not been submitted or is under consideration at any other journal.

## Acknowledgements

We would like to thank Rosa Ferrado for technical assistance and Lucía Garrido-Matilla, Shishir Baliyan, Natalia Puig, Ana Belén Sanz-Martos and María Roca for their help with animal procedures.

