## Supplementary Information for "Adolescent cannabinoid vapour exposure sex-dependently alters the relationship between vulnerability traits and ethanol self-administration and modifies naltrexone actions on ethanol intake in rats"

**SUPLEMENTARY MATERIAL**

**Supplementary Materials and Methods**

**Determination of CBD and THC in plasma by LC-MS/MS**

Chemicals, standards and materials

To develop the analytical method, the following solvents and chemical reagents were used: LC-MS grade acetonitrile, methanol, and formic acid (Carlo Erba Reagent, Val-de-Reuil, France); 99% isopropyl alcohol (Scharlab, Barcelona, Spain); and ammonium acetate and zinc sulphate (Merck, Darmstadt, Germany). Ultrapure water was obtained using a Milli-Q water purification system (Merck Millipore, Milford, MA, USA).

The standards (purity ≥ 98%) – tetrahydrocannabinol (THC), cannabidiol (CBD), and their internal standards (IS), THC-d₃ and CBD-d₃ – were purchased from Cayman Chemical (Michigan, USA). Commercial IS stock solutions (1,000 µg/mL) were prepared in methanol and stored in amber vials at −20 °C.

Solid-phase extraction (SPE) cartridges (Oasis HLB Prime; 60 mg/3 mL; hydrophilic-lipophilic balanced copolymer of divinylbenzene and N-vinylpyrrolidone) were purchased from Waters (Milford, MA, USA). The vacuum manifold used for SPE was supplied by Scharlab (Barcelona, Spain). An Eppendorf Centrifuge 5804 (Westbury, NY) capable of reaching 4,500 rpm, and a Biotage TurboVap LV sample evaporator (Charlotte, NC) operating with compressed nitrogen, were employed.

Sample treatment method

Sample treatment was performed via protein precipitation followed by SPE. A 500 µL plasma sample was spiked with 50 µL of each IS (0.1 µg/mL in 50:50 methanol/water; mix of CBD-d₃ and THC-d₃). Then, 125 µL of a 50:50 zinc sulphate/ammonium acetate solution (0.1 M) was added. Subsequently, 2 mL of ice-cold acetonitrile containing 0.1% formic acid was added dropwise, followed by vortexing for 40 seconds. Samples were centrifuged at 4,000 rpm for 5 minutes. The supernatant was transferred to a glass vial and diluted with 4 mL of water.

The extract was loaded onto an SPE cartridge by gravity. Cartridges were preconditioned with 3 mL of methanol followed by 3 mL of ultrapure water. After sample loading, the cartridge was washed with 2 mL of 25:75 (v/v) methanol/water and dried under vacuum (5 bar) for seven minutes. Elution was performed with 3 mL (two aliquots of 1.5 mL) of 90:10 (v/v) acetonitrile/isopropyl alcohol. The eluate was evaporated to dryness at 40 °C for 25 minutes under 0.2 Torr. Finally, the residue was reconstituted in 100 µL of acetonitrile followed by 100 µL of water, vortexed for two minutes, and transferred to an amber vial for LC-MS/MS analysis.

Analysis by LC-MS/MS

Analysis was performed using an LC-MS/MS system comprising an Agilent 1200 series HPLC (Palo Alto, CA, USA) coupled via an electrospray ionisation (ESI) interface (Agilent Technologies) to an Agilent 6410 triple quadrupole (QqQ) mass spectrometer. Chromatographic separation was conducted in positive ESI mode. A Kinetex F5 column (100 mm × 2.1 mm × 2.6 µm; Phenomenex, California, USA) was used. Eluent A consisted of 0.1% formic acid in ultrapure water, and eluent B of 0.1% formic acid in acetonitrile.

The gradient began with an isocratic elution of 70% eluent B for 4 minutes, followed by a linear increase to 98% over 2 minutes. This was held for 3 minutes, then reduced back to 70% over 60 seconds, followed by a 6-minute re-equilibration. Maximum pressure was set to 600 bar. The flow rate was 0.3 mL/min; the column oven was maintained at 50 °C, the autosampler at 4 °C, and the injection volume was 50 µL.

MS/MS conditions included: drying gas at 13 L/min at 350 °C, nebuliser pressure at 60 psi, capillary voltage at 3,500 V (positive polarity). Data acquisition was conducted in Multiple Reaction Monitoring (MRM) mode with a fragmentation voltage of 100 V. Compound-specific parameters are summarised in Table 1.

**Table 1.** LC-MS/MS parameters for cannabinoids in MRM mode: retention time (RT); precursor ion; and product ions (quantifier, Q, and qualifier, q) with their collision energies (CE) in parentheses.

| **Compounds** | **RT** | **Precursor ion**  **[MH^+^]** | **Product Ions** | |
| --- | --- | --- | --- | --- |
|  |  |  | **Q** | **q** |
| **CBD** | 3.2 min | 315.1 m/z | 193.1 m/z  (CE, 15 V) | 259.0 m/z  (CE, 20 V) |
| **THC** | 6.2 min | 315.1 m/z | 193.1 m/z  (CE, 20 V) | 259.0 m/z  (CE, 20 V) |
| **CBD-d^3^** | 3.2 min | 318.2 m/z | 196.0 m/z  (CE, 20 V) | 262.0 m/z  (CE, 20 V) |
| **THC-d^3^** | 6.2 min | 318.2 m/z | 196.0 m/z  (CE, 15 V) | 262.0 m/z  (CE, 15 V) |

The calibration method was performed in a plasma blank matrix with the addition of deuterated internal standards at each calibration point and for a linearity in the range of 1–100 ng/mL for THC and CBD. The LODs and LOQs were 0.3 ng/mL and 1 ng/mL for THC and CBD, respectively. The LODs and LOQs were 0.3 ng/mL and 1 ng/mL for THC and CBD, respectively.

**Operant oral ethanol self-administration procedure**

Pretraining

All experimental phases were conducted in operant conditioning chambers (Skinner boxes; 30 × 25 × 25 cm) specifically designed for behavioural testing. Each chamber was equipped with retractable levers, stimulus lights, food pellet dispensers, infrared beam sensors for head-entry detection into the food magazine, fans, ambient lighting, and grid floors capable of delivering electric footshocks (Med-Associates, USA). Experimental protocols were controlled using the Med-PC IV system. Ethanol was delivered via microlitre injection pumps (Harvard 22), which dispensed 100 μL of 15% (v/v) ethanol in tap water into a specially designed receptacle minimising evaporation.

Prior to initiating the ethanol self-administration protocol, rats underwent two preparatory phases: an autoshaping session (30 minutes) and an extended training session (60 minutes), both conducted in the same operant chambers. To promote fluid-seeking behaviour, animals were subjected to a 12-hour water deprivation period before each session.

During autoshaping, a single delivery of 100 μL of 15% (v/v) ethanol was made into the receptacle 3 minutes after session onset, followed by additional deliveries every 5 minutes. A stimulus light was illuminated for 4.5 seconds concurrently with each ethanol delivery. Levers were not present during this session.

Twenty-four hours later, rats underwent an extended training session lasting 60 minutes. Both active and inactive levers were extended. Each press on the active lever resulted in the delivery of 100 μL of 15% ethanol into the receptacle, accompanied by activation of both the stimulus light and the pump. The inactive lever had no programmed consequence and served as a control.

Following completion of these preparatory phases, the operant self-administration protocol commenced. From this point onward, rats had ad libitum access to tap water in their home cages. The protocol comprised seven consecutive phases, each designed to assess different behavioural dimensions of ethanol-seeking and consumption. Each phase consisted of multiple daily sessions (30 minutes each) conducted in the same operant chambers. During each session, both levers were extended. Responses on the active lever activated the pump, delivering 100 μL of 15% (v/v) ethanol into the receptacle, allowing voluntary oral consumption. The inactive lever remained non-functional, serving as a measure of non-specific motor activity.

**Supplementary Results**

**Hypothermia Induced by Adolescent THC vapour.**

This section provides a detailed description of the results obtained from the analysis of THC vapour-induced hypothermia. Specifically, it includes the outcomes of statistical analyses performed on the mean rectal temperature during the first two sessions (Pre and Post cannabinoid treatment) and the mean rectal temperature during the last two sessions (out of a total of nine sessions).

The results are reported for both within-subjects and between-subjects effects analyses, alongside multiple pairwise comparisons performed with Bonferroni corrections. Supplementary tables present all relevant statistical values, including F-statistics, p-values, partial eta-squared, confidence intervals, and observed power, to comprehensively support the findings detailed in the main text.

Mean rectal temperature of the first two sessions.

The measurement of rectal temperature was conducted to assess the physiological effects of cannabinoid exposure. We first analysed the averaged values of the first two vaporisation sessions (pre-vaporisation and post-vaporisation). Figure S1A, S1B, and S1C present the rectal temperature measurements recorded before (Pre) and after (Post) cannabinoid treatment administration. Figure S1A illustrates the overall mean rectal temperatures across the two initial sessions (Pre and Post) without separating by sex. Figures S1B and S1C provide a more detailed analysis, showing the mean rectal temperature changes separately for males (Figure S1B) and females (Figure S1C). These figures represent the averaged data from the first two sessions, highlighting the physiological effects of the treatment on rectal temperature.

A repeated-measures ANOVA was conducted to analyse the effects of vaporisation, sex, treatment, and their interactions on rectal temperature. A significant main effect of vaporisation was observed, F(_1, 70_) = 128.565, p < 0.001, ηp² = 0.647. The interaction between vaporisation x treatment was also significant, F(_1, 70_) = 28.899, p < 0.001, ηp² = 0.562. In contrast, the interaction between vaporisation x sex, F(_1, 70_) = 1.354, p = 0.249, ηp² = 0.019, was not statistically significant. Similarly, the interaction between vaporisation x sex x treatment was not significant, F(_1, 70_) = 1.454, p = 0.235, ηp² = 0.020 (See Table S1).

The analysis of the vaporisation x treatment interaction showed significant reductions in rectal temperature from pre- and post-vaporisation in the THC group (mean difference = 1.210, SE = 0.089, p < 0.001, 95% CI [1.033, 1.387]), THChighCBDlow group (mean difference = 0.328, SE = 0.089, p < 0.001, 95% CI [0.150, 0.505]) and in the THClowCBDhigh group (mean difference = 0.469, SE = 0.091, p < 0.001, 95% CI [0.288, 0.650]). In contrast, the VH control group did not show significant differences in rectal temperature between vaporisation and post-vaporisation (mean difference = 0.053, SE = 0.094, p = 0.580, 95% CI [-0.136, 0.241]). According to the mean differences reported, THC rats showed the most pronounced hypothermic effect among all treatment groups (Figure S1A and Table S2). We then examined differences among treatment groups post-vaporisation. The THC group had lower rectal temperature compared to the VH group (Mean difference = -1.094, SE = 0.143, p < 0.001, 95% CI [-1.481, -0.706]), confirming a pronounced hypothermic effect induced by THC. Additionally, when analysing the impact of co-administration of cannabinoids, significant differences were found between the THC and THChighCBDlow, which showed a significantly higher rectal temperature (Mean difference = 0.787, SE = 0.138, p < 0.001, 95% CI [0.411, 1.164]), suggesting that CBD co-administration at low doses mitigates the THC-induced hypothermic effect. Similarly, the combination of THClowCBDhigh ratio also had significantly higher temperatures than the THC (M = 0.662, SE = 0.140, p < 0.001, 95% CI [0.282, 1.042]), further indicating that CBD at high doses reduces hypothermic effect of THC. Finally, the treatment with THClowCBDhigh exhibited significantly lower temperatures compared to the Vehicle (VH) group (Mean difference = -0.431, SE = 0.144, p = 0.023, 95% CI [-0.823, -0.040]), suggesting that CBD at high doses may induce a mild hypothermic effect (Figure S1A and Table S3).

Our results demonstrate that THC exposure induces a significant hypothermic effect, as indicated by the marked reduction in rectal temperature following vaporization. However, co-administration of CBD attenuated this effect, in both low and high CBD doses (at the beginning of the treatment).

**Table S1.** Results of Within-Subjects Effects Analysis for Rectal Temperature Based on the Average of Pre-Treatment (Sessions 1 and 2) and Post-Treatment (Sessions 1 and 2) Measurements (Mean Transformed Variable).

| Source | | df | F-value | Sig.  (Significance) | Partial Eta Squared (η_p_²) | Observed Power^a^ |
| --- | --- | --- | --- | --- | --- | --- |
| Sessions | Greenhouse-Geisser | 1.000 | 128.565 | <.001 | .647 | 1.000 |
| Session*Sex | Greenhouse-Geisser | 1.000 | 1.354 | .249 | .019 | .209 |
| Sessions*Treatment | Greenhouse-Geisser | 3.000 | 29.898 | <.001 | .562 | 1.000 |
| Sessions*Sex*  Treatment | Greenhouse-Geisser | 3.000 | 1.454 | .235 | .059 | .369 |
| Error(Sessions) | Greenhouse-Geisser | 70.000 |  |  |  |  |

a. Calculated using alpha = .05

**Table S2.** Pairwise comparisons (treatment x vaporisation) (mean of sessions 1 and 2). The average of pre-treatment measurements and the post-treatment measurement were analysed (Mean Transformed Variable).

| Treatment | (I) Measurement – Pre. | (J) Measurement-Post. | Mean Difference  (I-J) | Std. Error | Sig.^b^ | 95% Confidence Interval^b^ | |
| --- | --- | --- | --- | --- | --- | --- | --- |
|  |  |  |  |  |  | Lower Bound | Lower Bound |
| VH | Pre | Post | .053 | .094 | .580 | -.136 | .241 |
|  | Post | Pre | -.053 | .094 | .580 | -.241 | .136 |
| THC | Pre | Post | 1.210^*^ | .089 | <.001 | 1.033 | 1.387 |
|  | Post | Pre | -1.210^*^ | .089 | <.001 | -1.387 | -1.033 |
| THChighCBDlow | Pre | Post | .328^*^ | .089 | <.001 | .150 | .505 |
|  | Post | Pre | -.328^*^ | .089 | <.001 | -.505 | -.150 |
| THClowCBDhigh | Pre | Post | .469^*^ | .091 | <.001 | .288 | .650 |
|  | Post | Pre | -.469^*^ | .091 | <.001 | -.650 | -.288 |
| Based on estimated marginal means | | | | | | | |
| *. The mean difference is significant at the .05 level. | | | | | | | |
| b. Adjustment for multiple comparisons: Bonferroni. | | | | | | | |

**Table S3.** Pairwise comparisons (vaporisation x treatment) (mean of sessions 1 and 2). The average of pre-treatment rectal measurements and the post-treatment measurement were analysed (Mean Transformed Variable).

| Sessions | (I) Treatment | (J) Treatment | Mean Difference  (I-J) | Std. Error | Sig.^b^ | 95% Confidence Interval^b^ | |
| --- | --- | --- | --- | --- | --- | --- | --- |
|  |  |  |  |  |  | Lower Bound | Lower Bound |
| Pre- | VH | THC | -.064 | .054 | 1.000 | -.211 | .083 |
|  |  | THChighCBDlow | .031 | .054 | 1.000 | -.116 | .178 |
|  |  | THClowCBDhigh | .015 | .055 | 1.000 | -.133 | .164 |
|  | THC | VH | .064 | .054 | 1.000 | -.083 | .211 |
|  |  | THChighCBDlow | .095 | .053 | .449 | -.048 | .238 |
|  |  | THClowCBDhigh | .079 | .053 | .848 | -.065 | .223 |
|  | THChighCBDlow | VH | -.031 | .054 | 1.000 | -.178 | .116 |
|  |  | THC | -.095 | .053 | .449 | -.238 | .048 |
|  |  | THClowCBDhigh | -.016 | .053 | 1.000 | -.160 | .128 |
|  | THClowCBDhigh | VH | -.015 | .055 | 1.000 | -.164 | .133 |
|  |  | THC | -.079 | .053 | .848 | -.223 | .065 |
|  |  | THChighCBDlow | .016 | .053 | 1.000 | -.128 | .160 |
| Post- | VH | THC | 1.094^*^ | .143 | ***<.001*** | .706 | 1.481 |
|  |  | THChighCBDlow | .306 | .143 | .212 | -.081 | .694 |
|  |  | THClowCBDhigh | .431^*^ | .144 | ***.023*** | .040 | .823 |
|  | THC | VH | -1.094^*^ | .143 | ***<.001*** | -1.481 | -.706 |
|  |  | THChighCBDlow | -.787^*^ | .138 | ***<.001*** | -1.164 | -.411 |
|  |  | THClowCBDhigh | -.662^*^ | .140 | ***<.001*** | -1.042 | -.282 |
|  | THChighCBDlow | VH | -.306 | .143 | .212 | -.694 | .081 |
|  |  | THC | .787^*^ | .138 | ***<.001*** | .411 | 1.164 |
|  |  | THClowCBDhigh | .125 | .140 | 1.000 | -.255 | .505 |
|  | THClowCBDhigh | VH | -.431^*^ | .144 | ***.023*** | -.823 | -.040 |
|  |  | THC | .662^*^ | .140 | ***<.001*** | .282 | 1.042 |
|  |  | THChighCBDlow | -.125 | .140 | 1.000 | -.505 | .255 |
| Based on estimated marginal means | | | | | | | |
| *. The mean difference is significant at the .05 level. | | | | | | | |
| b. Adjustment for multiple comparisons: Bonferroni. | | | | | | | |

Mean rectal temperature of the last two sessions.

We then looked at the mean rectal temperatures from the last two treatment sessions (pre- and post-vaporisation). Figure S1D showing the overall mean and Figures S1E and S1F detailing the results for males and females, respectively.

A significant main effect of vaporisation was observed, F(_1,70_) = 80.817, p < 0.001, ηp² = 0.536, indicating substantial changes in rectal temperature. The interaction between vaporisation x treatment was also significant, F(_3,70_) = 22.714, p < 0.001, ηp² = 0.493, revealing that the effect of vaporisation on rectal temperature depended on the treatment condition. In contrast, the interaction between vaporisation x sex was not significant, (F(_1,70_) = 1.893, p = 0.173, ηp² = 0.026). Similarly, the three-way interaction between vaporisation x sex x treatment was not significant, F(_3,70_) = 1.877, p = 0.141, ηp² = 0.074.

The analysis of the vaporisation x treatment interaction revealed significant temperature reductions in the THC group, indicating a hypothermic effect (mean difference = 1.000, SE = 0.097, p < 0.001, 95% CI [0.806, 1.194]). Similarly, significant reductions were observed in THChighCBDlow group which displayed a significant decrease in rectal temperature post-vaporisation (mean difference = 0.653, SE = 0.097, p < 0.001, 95% CI [0.458, 0.847]). The THClowCBDhigh showed a smaller but still significant reduction (Mean difference = 0.217, SE = 0.099, p = 0.033, 95% CI [0.018, 0.415]). In contrast, the VH group did not show significant differences in rectal temperature (mean difference = -0.082, SE = 0.103, p = 0.431, 95% CI [-0.288, 0.124]) (Figure S1B and Table S5).

The then analysed group differences post-vaporisation. Pairwise comparisons of treatment effects post-vaporisation revealed significant differences in rectal temperature between treatment groups. The THC group showed a significantly lower rectal temperature compared to the Vehicle (VH) group (Mean difference = -1.113, SE = 0.139, p < 0.001, 95% CI [-1.492, -0.734]), confirming a strong hypothermic effect induced by THC. The THChighCBDlow also exhibited significantly lower rectal temperatures compared to the Vehicle (VH) group (Mean difference = -0.768, SE = 0.139, p < 0.001, 95% CI [-1.147, -0.389]), suggesting that this combination maintains a hypothermic effect. Similarly, the THClowCBDhigh had significantly lower temperatures compared to Vehicle (VH) (M = -0.425, SE = 0.141, p = 0.021, 95% CI [-0.807, -0.042]), indicating a mild but significant hypothermic effect. When comparing THC vs. THClowCBDhigh, the THC showed significantly lower temperatures (Mean difference = -0.689, SE = 0.137, p < 0.001, 95% CI [-1.060, -0.317]), suggesting that the high CBD dose reduced the THC-induced hypothermia. In contrast, the comparison between THC and THChighCBDlow did not reach statistical significance (Mean difference = -0.345, SE = 0.135, p = 0.078, 95% CI [-0.712, 0.022]), suggesting that CBD at a lower dose had a weaker effect in counteracting THC hypothermic actions. (Figure S1B and Table S6).

**Table S4.** Results of Within-Subjects Effects Analysis for Rectal Temperature Based on the Average of Pre-Treatment (Sessions 8 and 9) and Post-Treatment (Sessions 8 and 9) Measurements (Mean Transformed Variable).

| Source | | df | F-value | Sig.  (Significance) | Partial Eta Squared (η_p_²) | Observed Power^a^ |
| --- | --- | --- | --- | --- | --- | --- |
| Sessions | Greenhouse-Geisser | 1.000 | 80.817 | ***<.001*** | .536 | 1.000 |
| Session*Sex | Greenhouse-Geisser | 1.000 | 1.893 | .173 | .026 | .274 |
| Sessions*Treatment | Greenhouse-Geisser | 3.000 | 22.714 | ***<.001*** | .493 | 1.000 |
| Sessions*Sex*  Treatment | Greenhouse-Geisser | 3.000 | 1.877 | .141 | .074 | .466 |
| Error(Sessions) | Greenhouse-Geisser | 70.000 |  |  |  |  |

a. Calculated using alpha = .05

**Table S5.** Pairwise comparisons (treatment x vaporisation) (mean of sessions 8 and 9). The average of pre-treatment rectal measurements and the post-treatment measurement were analysed (Mean Transformed Variable).

| Treatment | (I) Sessions – Pre. | (J) Sessions-Post. | Mean Difference  (I-J) | Std. Error | Sig.^b^ | 95% Confidence Interval^b^ | |
| --- | --- | --- | --- | --- | --- | --- | --- |
|  |  |  |  |  |  | Lower Bound | Lower Bound |
| VH | Pre | Post | -.082 | .103 | .431 | -.288 | .124 |
|  | Post | Pre | .082 | .103 | .431 | -.124 | .288 |
| THC | Pre | Post | 1.000 | .097 | ***<.001*** | .806 | 1.194 |
|  | Post | Pre | -1.000 | .097 | ***<.001*** | -1.194 | -.806 |
| THChighCBDlow | Pre | Post | .653 | .097 | ***<.001*** | .458 | .847 |
|  | Post | Pre | -.653 | .097 | ***<.001*** | -.847 | -.458 |
| THClowCBDhigh | Pre | Post | .217 | .099 | ***.033*** | .018 | .415 |
|  | Post | Pre | -.217 | .099 | ***.033*** | -.415 | -.018 |
| Based on estimated marginal means | | | | | | | |
| *. The mean difference is significant at the .05 level. | | | | | | | |
| b. Adjustment for multiple comparisons: Bonferroni. | | | | | | | |

**Table S6.** Pairwise comparisons (vaporisation x treatment) (mean of sessions 8 and 9). The average of pre-treatment rectal measurements and the post-treatment measurement were analysed (Mean Transformed Variable).

| Sessions | (I) Treatment | (J) Treatment | Mean Difference  (I-J) | Std. Error | Sig.^b^ | 95% Confidence Interval^b^ | |
| --- | --- | --- | --- | --- | --- | --- | --- |
|  |  |  |  |  |  | Lower Bound | Lower Bound |
| Pre- | VH | THC | .031 | .063 | 1.000 | -.141 | .203 |
|  |  | THChighCBDlow | .034 | .063 | 1.000 | -.138 | .206 |
|  |  | THClowCBDhigh | .126 | .064 | .316 | -.048 | .300 |
|  | THC | VH | -.031 | .063 | 1.000 | -.203 | .141 |
|  |  | THChighCBDlow | .002 | .061 | 1.000 | -.164 | .169 |
|  |  | THClowCBDhigh | .095 | .062 | .788 | -.074 | .263 |
|  | THChighCBDlow | VH | -.034 | .063 | 1.000 | -.206 | .138 |
|  |  | THC | -.002 | .061 | 1.000 | -.169 | .164 |
|  |  | THClowCBDhigh | .092 | .062 | .850 | -.076 | .261 |
|  | THClowCBDhigh | VH | -.126 | .064 | .316 | -.300 | .048 |
|  |  | THC | -.095 | .062 | .788 | -.263 | .074 |
|  |  | THChighCBDlow | -.092 | .062 | .850 | -.261 | .076 |
| Post- | VH | THC | 1.113^*^ | .139 | ***<.001*** | .734 | 1.492 |
|  |  | THChighCBDlow | .768^*^ | .139 | ***<.001*** | .389 | 1.147 |
|  |  | THClowCBDhigh | .425^*^ | .141 | ***.021*** | .042 | .807 |
|  | THC | VH | -1.113^*^ | .139 | ***<.001*** | -1.492 | -.734 |
|  |  | THChighCBDlow | -.345 | .135 | .078 | -.712 | .022 |
|  |  | THClowCBDhigh | -.689^*^ | .137 | ***<.001*** | -1.060 | -.317 |
|  | THChighCBDlow | VH | -.768^*^ | .139 | ***<.001*** | -1.147 | -.389 |
|  |  | THC | .345 | .135 | .078 | -.022 | .712 |
|  |  | THClowCBDhigh | -.344 | .137 | .086 | -.715 | .028 |
|  | THClowCBDhigh | VH | -.425^*^ | .141 | ***.021*** | -.807 | -.042 |
|  |  | THC | .689^*^ | .137 | ***<.001*** | .317 | 1.060 |
|  |  | THChighCBDlow | .344 | .137 | .086 | -.028 | .715 |
| Based on estimated marginal means | | | | | | | |
| *. The mean difference is significant at the .05 level. | | | | | | | |
| b. Adjustment for multiple comparisons: Bonferroni. | | | | | | | |

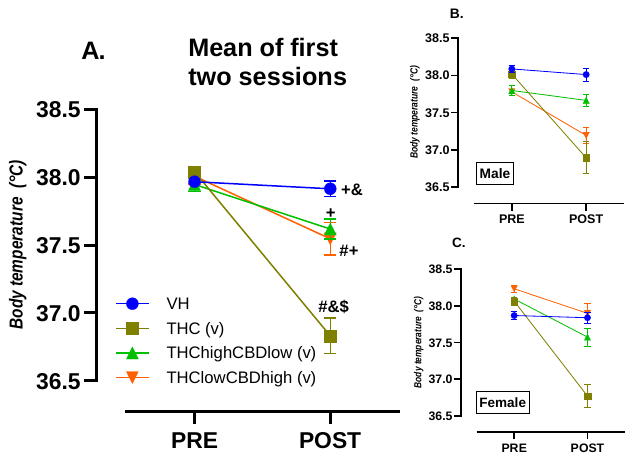

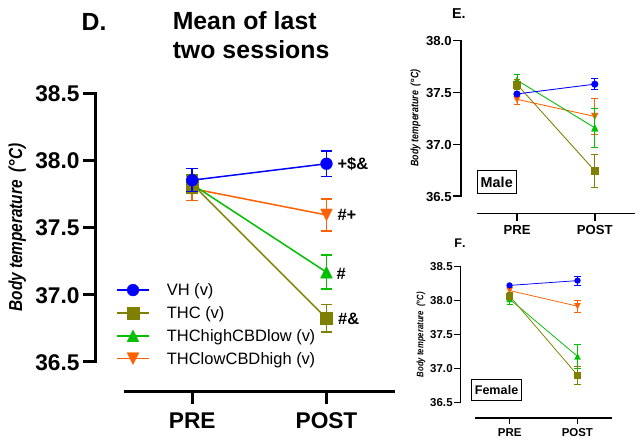

**Figure S1. Effects of cannabinoid vaporization on rectal temperature.** (A) Post-treatment differences in rectal temperature were observed, with a significant interaction between vaporisation × treatment (F(3, 70) = 29.898, p < 0.001, ηp² p = 0.562). This data represents the mean rectal temperature from the first two sessions. (B, C) Post-vaporisation, differences in rectal temperature analysed by the interaction between sex × vaporisation were not significant (F(1, 70) = 1.354, p = 0.249, ηp² p = 0.019). (D) Post-vaporisation, differences in rectal temperature were observed with a significant interaction between vaporisation × treatment (F(3, 70) = 22.714, p < 0.001, ηp² p = 0.493). This data represents the mean rectal temperature from the last two sessions. (E, F) Post-vaporisation, differences in rectal temperature analysed by the interaction between sex × vaporisation were not significant (F(1, 70) = 1.893, p = 0.173, ηp² p = 0.026). All data are expressed as mean ± S.E.M. Symbols denote significant differences (*p < 0.05), from the vehicle (VH) is indicated by ^#^, THC is indicated by ^+^, THChighCBDlow is indicated by ^$^ and THClowCBDhigh is indicated by ^&^. Significant effect of the within-subjects factor (pre-post vaporisation effect of rectal temperature) is denoted by ‘v’. All data are expressed as mean ± S.E.M.

**Behavioural Assessment**

Saccharine preference Test

Saccharine preference developed over time (significant effect of the sessions factor (F_6.682, 467.726_=36.865 p<0.0001). We found significant differences in saccharine preference due to the interaction of the factors sessions x treatment (F_(20.045, 630)_ = 2.079, p < 0.004, η_p_² = 0.082) but there were group differences only in sessions 2 and 10. There was a trend for an effect of the sessions x sex (F_(6.682, 630)_ = 2.003, p < 0.056, η_p_² = 0.028) interaction or sessions x sex x treatment (F_(20.045, 630)_ = 1.027, p < 0.428, η_p_² = 0.042) interactions. In addition, there were no effects of the main factor sex and treatment. (Figure S2).

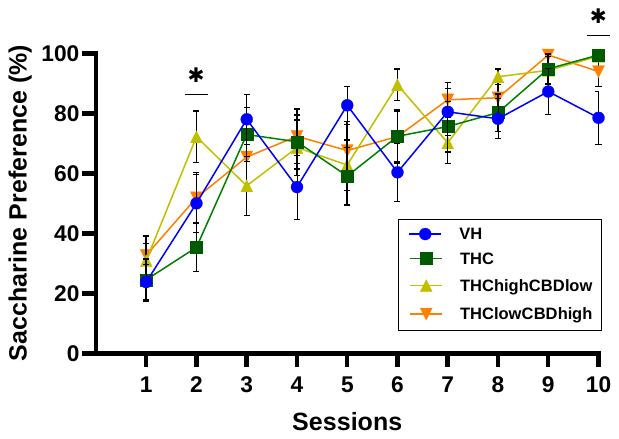

**Figure S2. Saccharine preference test.** he graph shows the percentage of saccharin intake relative to pure water across sessions (± S.E.M.). Overall, all rats exhibited a high preference for saccharin-containing water. Neither treatment nor sex affected anhedonia behaviour. Significance symbols are defined as follows: * p < 0.05.

Acquisition of ethanol self-administration (FR1 and FR2).

During the FR1 phase, animals progressively learned to press the active lever to obtain a dose of ethanol, as evidenced by a significant main effect of sessions (F_(13.026, 898.771)_ = 4.837, p < 0.001, ηp² = 0.066), lever (F_(1, 69)_ = 93.425, p < 0.001, ηp² = 0.575) and by the interaction of the sessions x lever (F_(11.812, 815.001)_ = 16.107, p < 0.001, ηp² = 0.189), This result confirms that ethanol self-administration behaviour increased over time as sessions progressed (Figure S3A). The interaction between sessions x treatment was also significant (F_(39.077, 898.771)_ = 1.606, p = 0.012, ηp² = 0.065), suggesting that acquisition rates varied between treatment groups. Post-hoc comparisons using Bonferroni corrections revealed that animals treated with THC pressed the active lever fewer times compared to the VH group and other treatment groups across the sessions of FR1. In session 1, the mean difference between THC and VH was -11.497 (SE = 3.362, p = 0.006, 95% CI [-20.630, -2.365]). Similarly, the THChighCBDlow group also exhibited fewer active lever presses compared to the VH group in session 1 (mean difference = -12.175, SE = 3.319, p = 0.003, 95% CI [-21.191, -3.159]). The THClowCBDhigh group showed a similar trend, with significantly fewer active lever presses than the VH group during session 1 (mean difference = -10.075, SE = 3.319, p = 0.020, 95% CI [-19.091, -1.059]). This trend only persisted in session 5, when the THC group pressed the active lever significantly fewer times than the VH group (mean difference = -6.301, SE = 2.211, p = 0.035, 95% CI [-12.308, -0.294]). Additionally, the THClowCBDhigh group demonstrated significantly lower active lever presses compared to the VH group in session 5 (mean difference = -8.212, SE = 2.183, p = 0.002, 95% CI [-14.142, -2.283]). However, no significant differences were observed between THC and the combination treatment groups (THChighCBDlow or THClowCBDhigh) during any session (p > 0.05). In considering the global context of all the acquisition sessions, these effects appear to be spurious, as no significant differences were observed in other sessions.

We also observed a main effect of sex, with females exhibiting significantly more lever presses compared to males (F_(1, 69)_ = 14.138, p < 0.001, ηp² = 0.170). However, no significant interaction effects were found for sessions x sex (F_(13.026, 898.771)_ = 0.401, p = 0.970, ηp² = 0.006) or sessions x sex x treatment (F_(39.077, 898.771)_ = 1.003, p = 0.467, ηp² = 0.042), indicating that neither sex nor its interaction with treatment influenced acquisition rates during FR1.

The interaction between lever x sex was not statistically significant (F_(1, 69)_ = 3.473, p = 0.067, ηp² = 0.048). Similarly, the interaction between lever x treatment was not significant (F_(3, 69)_ = 0.308, p = 0.819, ηp² = 0.013), nor was the three-way interaction of lever x sex x treatment (F_(3, 69)_ = 1.581, p = 0.202, ηp² = 0.064). These findings indicate that the preference for the active lever over the inactive lever was consistent across groups and sexes and not influenced by treatment. The interaction between sessions x lever x treatment was not statistically significant (F_(35.435, 898.771)_ = 1.013, p = 0.449, ηp² = 0.042), nor was the four-way interaction between Sessions x Lever x Sex x Treatment (F_(35.435, 898.771)_ = 1.260, p = 0.145, ηp² = 0.052).

Lastly, analysis of the mean number of active lever presses during the last three sessions revealed a statistically significant between-subjects effect for the sex factor (F_(1, 69)_ = 4.567, p = 0.036, ηp² = 0.062). Post-hoc comparisons with Bonferroni corrections showed that males pressed the active lever less frequently than females (mean difference = -4.707, SE = 2.203, p = 0.036, 95% CI [-9.101, -0.313]). Significant effects by treatment were not observed (F_(3, 69)_ = 0.605, p = 0.617, ηp² = 0.026) and also by the interactions of Sex x treatment (F_(3, 69)_ = 0.787, p = 0.505, ηp² = 0.033).

These results suggest that animals treated with cannabinoid vapor during adolescence did not show a different operant behaviour for ethanol self-administration compared to the VH group.

During the FR2 phase, a significant main effect of lever was observed (F_(1, 69)_ = 277.028, p < 0.001, ηp² = 0.801), indicating a strong preference for the active lever over the inactive lever, and thus demonstrating that the animals had successfully acquired the operant behaviour. Additionally, a significant interaction between lever and sex (F_(1, 69)_ = 5.339, p = 0.024, ηp² = 0.072) suggested that females exhibited slightly higher active lever-pressing activity compared to males, showing a marked sexual difference. However, there was no significant interaction between Lever x treatment (F_(3, 69)_ = 0.718, p = 0.544, ηp² = 0.030) or the three-way interaction of lever x sex x group (F_(3, 69)_ = 1.075, p = 0.366, ηp² = 0.045). A significant main effect of the sessions was found (F_(13, 897)_ = 7.849, p < 0.001, ηp² = 0.102), demonstrating progressive learning over time. However, neither the sessions x sex interaction (F_(13, 897)_ = 0.921, p = 0.530, ηp² = 0.013), nor sessions x group interaction (F_(39, 897)_ = 1.244, p = 0.148, ηp² = 0.051), nor the sessions x sex x group three-way interaction (F_(39, 897)_ = 0.948, p = 0.563, ηp² = 0.040) reached significance. Interestingly, the interaction between lever x sessions was significant (F_(13, 897)_ = 9.734, p < 0.001, ηp² = 0.124), showing that lever preference strengthened across sessions as the animals learned the operant task and to discriminate the active lever from the inactive lever. However, no significant interactions were found between lever x sessions x sex (F_(13, 897)_ = 0.956, p = 0.494, ηp² = 0.014), or between lever x sessions x group (F_(39, 897)_ = 0.944, p = 0.569, ηp² = 0.039), or the four-way interaction when all factors were combined (F_(39, 897)_ = 0.982, p = 0.503, ηp² = 0.041).

The analysis of the mean number of active lever presses during the last three sessions revealed a statistically significant between-subjects effect for the sex factor (F_(1, 69)_ = 6.874, p = 0.011, ηp² = 0.091). Post-hoc comparisons with Bonferroni corrections showed that males pressed the active lever less frequently than females (mean difference = -8.949, SE = 3.413, p = 0.011, 95% CI [-15.757, -2.140]). Significant effects by treatment were not observed (F_(3, 69)_ = 0849, p = 0.472, ηp² = 0.036) as neither were sex x treatment interactions (F_(3, 69)_ = 0.797, p = 0.500, ηp² = 0.033).

Subsequently, when animals transitioned to the ethanol acquisition phase under a fixed ratio 2 schedule (FR2, i.e., two responses), no significant interaction was found. (Figure S3B). Thus, all animals, regardless of treatment condition, successfully acquired the discrimination between the active and inactive lever for ethanol self-administration.

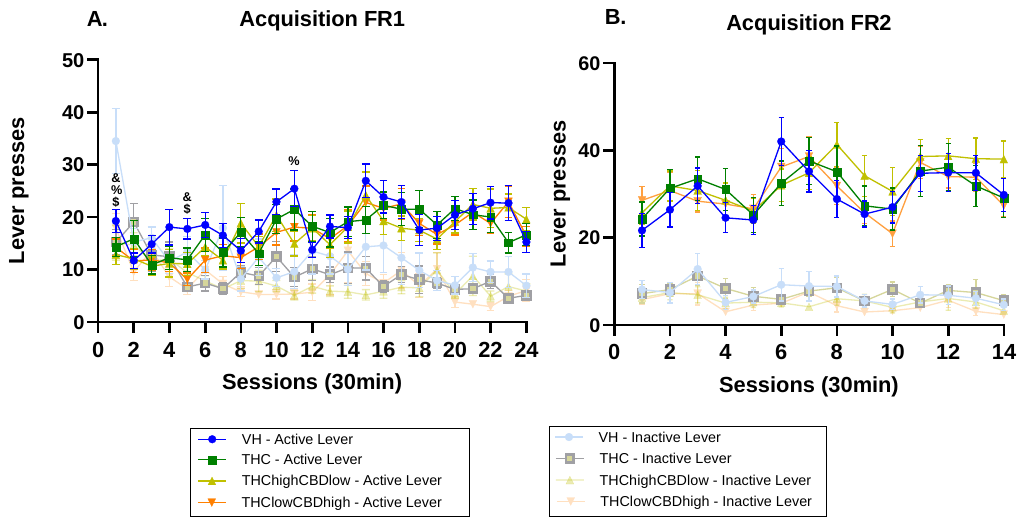

**Figure S3**. Active lever presses across sessions in the different treatment groups under FR1. (A). Data are presented as mean ± S.E.M. Symbols represent differences between treatment groups. Significant differences were observed in Sessions 1, 5, and 11. (B). Active lever presses under the fixed ratio 2 (FR2) schedule across sessions during the ethanol acquisition phase. Data are presented as mean ± S.E.M. No significant interaction was observed between Sessions and Treatment group. All animals, irrespective of treatment condition, acquired the discrimination between the active and inactive lever required for ethanol self-administration. Specifically, significant differences between VH and THC are indicated by $, between VH and THChighCBDlow by %, and between VH and THClowCBDhigh by &, according to Bonferroni-corrected post hoc tests. Statistical significance was set at p < 0.05.

Progressive ratio

A mixed-design ANOVA revealed a significant main effect of Lever, (F_(1, 69)_ = 126.392, p < 0.001, ηp² = 0.647). No significant interactions were found between Lever x Sex, (F_(1, 69)_ = 1.029, p = 0.314, ηp² = 0.015), or between Lever and Treatment, (F_(3, 69)_ = 1.177, p = 0.325, ηp² = 0.049) (Figure S4). Additionally, the three-way interaction of Lever × Sex × Treatment was not significant, (F_(3, 69)_ = 0.394, p = 0.757, ηp² = 0.017). Pairwise comparisons with Bonferroni adjustment confirmed that all animals, regardless of group or sex, exhibited a higher number of responses on the active lever than on the inactive lever (mean difference = 37.308, p < 0.001). Neither sex nor treatment had any significant impact on operant behaviour related to the motivation to consume ethanol.

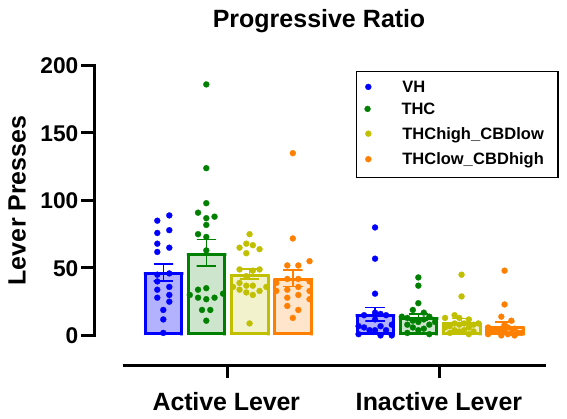

**Figure S4.** Number of lever presses on the active and inactive levers during the progressive ratio schedule, used to assess motivation for ethanol self-administration across treatment groups. Animals from all groups (VH, THC, THChighCBDlow, THClowCBDhigh) showed a significantly higher number of responses on the active lever compared to the inactive lever. No significant differences were observed between treatment groups or sexes. Data are expressed as mean ± SEM.

Stabilization phase after progressive ratio

Following the assessment of ethanol consumption motivation through the progressive ratio test, animals were re-exposed to a stabilization phase under a fixed ratio 2 (FR2) reinforcement schedule. A univariate ANOVA conducted on the average number of lever presses across the last three sessions revealed a significant main effect of Sex on overall lever-pressing behaviour (F_(1, 69)_ = 8.193, p = 0.006, ηp² = 0.106). Neither Treatment (F_(3, 69)_ = 0.418, p = 0.741, ηp² = 0.018) nor the Sex × Treatment interaction (F(3, 69) = 0.270, p = 0.847, ηp² = 0.012) reached significance. Bonferroni-adjusted pairwise comparisons showed that females made significantly more active lever presses compared to males during this stabilization phase (Mean difference = 7.262, SE = 2.537, p = 0.006, 95% CI [2.201, 12.323]) (Figure S5).

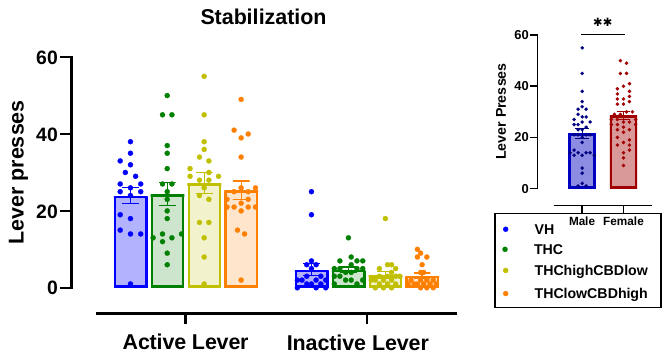

**Figure S5.** Lever pressing behaviour across treatment groups and sexes during stabilization phase. The main graph shows the mean (± SEM) number of lever presses on the active and inactive levers during the last three sessions of the stabilization phase under a fixed-ratio 2 (FR2) schedule. Across all groups, rats consistently pressed more on the active than the inactive lever. The inset plot on the right displays the mean active lever presses separated by sex. Females exhibited significantly higher active lever responding than males (p = 0.006), indicating greater ethanol-seeking behaviour during the stabilization phase. (* *p* < 0.01).

Analysis of Compulsive Ethanol Intake and Operant Behaviour.

During this phase, the behaviour of the rats was studied in an operant self-administration paradigm for ethanol. Here, animals were again subjected to a fixed ratio 2 (FR2) schedule, but pressing the active lever first delivered a 0.2 mA footshock to punish the seeking behaviour, followed by activation of the pump that dispensed a dose of ethanol, which the rat then consumed. Pearson correlation analyses indicated a significant positive association between the number of lever presses, reinforcers obtained, and head entries into the drinking area during the reinforcement period, confirming that the rats were actively consuming the ethanol (Figure S6A and B and Table S7).

Significant differences were observed for the main effect of lever (F_(1, 67)_ = 262.079, p < 0.001, ηp² = 0.796), confirming a strong preference for the active lever despite the punishment. Additionally, a significant interaction between lever and sex factors (F_(1, 67)_ = 40.474, p < 0.001, ηp² = 0.377). Post-hoc comparisons with Bonferroni corrections showed that males pressed the active lever less frequently than females (mean difference = -12.920, SE = 1.864, p < 0.001, 95% CI [-16.669, -9.230]).

Also, the analysis of lever x sessions revealed a significant effect (F_(11.468, 768.338)_ = 13.080, p < 0.001, ηp² = 0.163). Post-hoc comparisons with Bonferroni corrections showed that active lever presses were higher than inactive lever presses across all sessions. The three-way interaction of lever x sessions x sex showed significant effect (F_(11.468, 768.338)_ = 4.343, p < 0.001, ηp² = 0.061) (Figure 3D). Post-hoc comparisons with Bonferroni corrections showed that males and females press the active lever to a similar level during the sessions 1 and 2, but from session 3 onward, females pressed more the active lever than males across the remaining sessions.

The analysis of the sessions factor also revealed a significant effect (F_(11.260, 754.392)_ = 16.840, p < 0.001, ηp² = 0.201), indicating progressive changes in lever pressing over time. A significant interaction between sessions x sex was observed (F_(11.260, 754.392)_ = 5.253, p < 0.001, ηp² = 0.073). Post-hoc comparisons with Bonferroni corrections showed negative mean differences of lever pressing compared to females across all sessions, indicating that lever presses of the males were consistently lower than those of females in each session.

There was no significant effect of the lever x sessions x treatment interaction (F_(34.403, 768.338)_ = 1.040, p = 0.407, ηp² = 0.045) and neither there were of the lever x sessions x sex x treatment interaction (F_(34.403, 768.338)_ = 1.017, p = 0.442, ηp² = 0.044).

Lastly, there were no significant effects found for the interaction between the factors sessions x treatment (F_(33.779, 754.392)_ = 0.925, p = 0.592, ηp² = 0.040) and neither between the interaction of the factors sessions x sex x treatment presented a statistical effect (F_(33.779, 754.392)_ = 0.891, p = 0.648, ηp² = 0.038).

We also calculated the percentage decrease in the operant response (active lever presses) after the onset of electric shocks as punishment, relative to the average of the last three sessions of the stabilization phase (without electric shocks). After session 4, the rats reached their maximum decrease, and from session 5 to 7, they achieved their peak recovery. We observed significant differences due to the interaction between the factors sessions x sex (F_(1, 67)_ = 4.764, P = 0.033, ηp² = 0.066) (Figure 3E), with females exhibiting more pronounced compulsive behaviour, compared to males, in terms of ethanol self-administration. We did not observe significant differences between the factor session x treatment (F(3.000, 67.000) = 1.225, P = 0.307, ηp² = 0.052) or for the interaction between the factors sessions x sex x treatment (F(3.000, 67.000) = 0.539, P = 0.657, ηp² = 0.024).

Additionally, the between-subjects analysis of variance (ANOVA) revealed a significant effect of sex on the number of reward obtained (F_(1,68)_ = 30.049, p < 0.001, ηp² = 0.306), the number of lever presses (F_(1,68)_ = 30.960, p < 0.001, ηp² = 0.313), and head entries during the reinforcement period (F_(1,68)_ = 26.657, p < 0.001, ηp² = 0.282). Pairwise comparisons indicated that females received more reinforcers than males (7.112, p < 0.001), performed more lever presses (14.445, p < 0.001), and showed more head entries during the reinforcement period (6.205, p < 0.001). No significant effects were found by the interaction of treatment x rewards (F_(3,68)_ = 0.637, p < 0.594, ηp² = 0.027), treatment x lever presses (F_(3,68)_ = 0.624, p < 0.602, ηp² = 0.027) and treatment x head entries (F_(3,68)_ = 0.177, p < 0.161, ηp² = 0.072).

Also, interactions between sex x treatment x reward (F_(3,68)_ = 1.068, p < 0.369, ηp² = 0.045), sex x treatment x lever presses (F_(3,68)_ = 1.054, p < 0.374, ηp² = 0.044) and sex x treatment x head entries (F_(3,68)_ = 0.590, p < 0.624, ηp² = 0.025) were not found to be significant, suggesting that cannabis exposure during adolescence did not alter ethanol self-administration in adulthood.

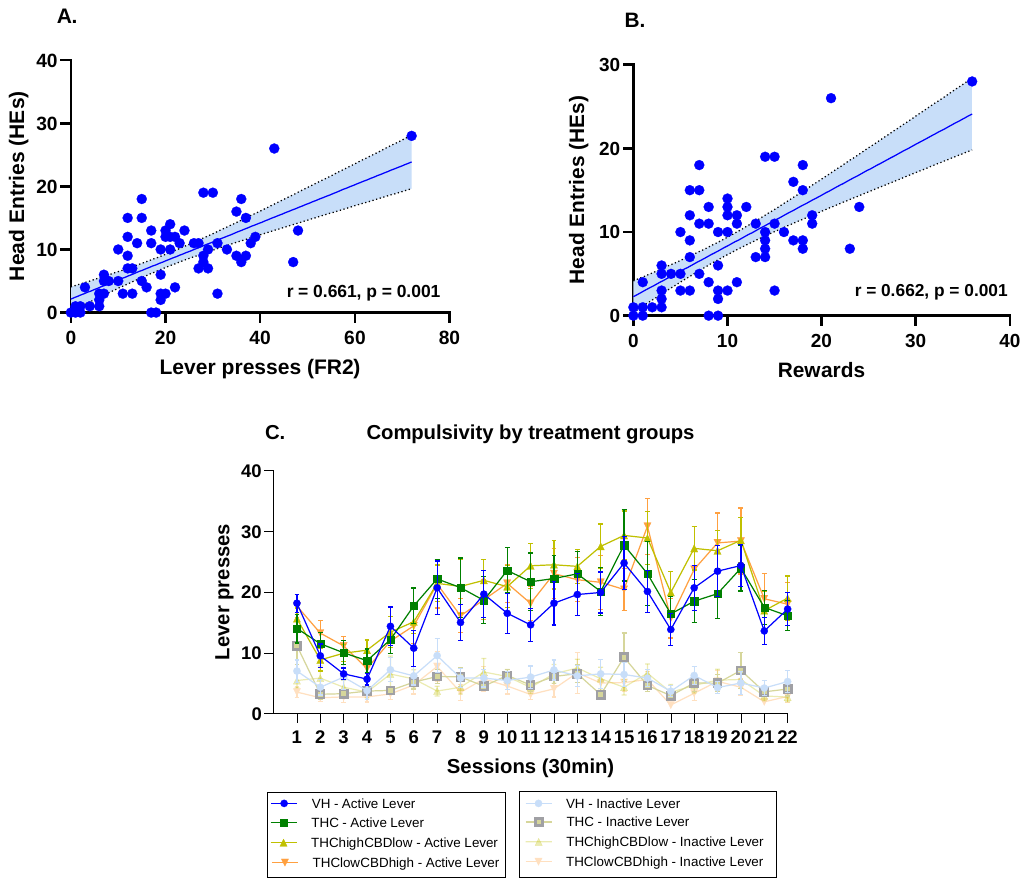

**Figure S6. Pearson Correlations Between Operant Responses, Reinforcement, and Ethanol-Seeking Behaviour.** (A) A significant correlation between head entries and lever presses suggests that operant responses are linked to ethanol-seeking behaviour, rather than random exploratory activity. (± S.E.M.). (B) A significant positive correlation was found between rewards obtained and head entries during the reinforcement period (± S.E.M.), indicating that ethanol delivery is associated with increased approach behaviour to the drinking area. (C) Active and inactive lever pressing during the compulsivity test across treatment groups (± S.E.M.). All treatment groups showed greater responding on the active lever compared to the inactive one, indicating persistence of ethanol-seeking despite punishment. Shaded lines represent inactive lever.

**Table S7**. Pearson correlation between variables during compulsive seeking phase

| Variable | Variable | Pearson correlation ( r ) | Sig. (Significance) |
| --- | --- | --- | --- |
| Lever presses (LP1) | Rewards | 0.999 | 0.001 |
| Rewards | Head Entries (HES) | 0.662 | 0.001 |
| Head Entries (HEs) | Lever Presses (LP1) | 0.661 | 0.001 |

**Note.** The values of Head Entries correspond to those recorded after pressing the lever, during the period when the pump administered the ethanol dose.

Correlation between behavioural characterization and ethanol self-administration phase.

Pearson correlation analyses were performed to explore the predictive value of different behavioural parameters on ethanol consumption and compulsive drinking in adulthood. The analyses revealed a total of six significant correlations in females and only one significant correlation in males. The table 1 below summarizes the significant correlations detected in both sexes.

**Table S8.** Significant Pearson correlations between behavioural parameters and ethanol consumption or compulsivity, separated by sex.

| Variable | Variable | Sex | Treatment | Pearson correlation ( r ) | Sig. (Significance) |
| --- | --- | --- | --- | --- | --- |
| SP | COMP | M | THC | -0.77 | 0.015 |
| EPM | LP1_FR1 | F | VH | 0.64 | 0.047 |
| NP | LP1_FR2 | F | VH | 0.77 | 0.010 |
| PCAIndex | BP | F | THC | -0.72 | 0.020 |
| PCAIndex | COMP | F | THC | -0.88 | 0.004 |
| NP | LP1_FR1 | F | THChighCBDlow | 0.72 | 0.018 |
| OF | BP | F | THClowCBDhigh | 0.69 | 0.030 |

**Abbreviations:** SP = Saccharine Preference; EPM = Elevated Plus Maze; NP = Novelty Preference; OF = Open Field; PCAIndex = Sign/Goal Tracking Bias; LP1_FR1 = Active Lever Presses under Fixed Ratio 1 (FR1); LP1_FR2 = Active Lever Presses under Fixed Ratio 2 (FR2); Break-Point = Maximum ratio completed during progressive ratio testing; COMP = Compulsivity (i.e., percentage of lever presses maintained despite foot shock punishment).
